# Continuous Flash Suppression responses in mouse visual cortex: stimulus laterality and anesthesia effects

**DOI:** 10.1101/2025.11.05.686768

**Authors:** Mathis Bassler, Lilian Emming, Christopher J. Whyte, Gerjan Huis In ‘t Veld, Mototaka Suzuki, Cyriel M. A. Pennartz

## Abstract

We investigated whether binocularly conflicting stimuli adapted from primate binocular rivalry studies could induce binocular response suppression in mouse visual cortex. We presented binocularly conflicting stimuli adapted from the paradigm of continuous flash suppression to awake and anesthetized mice and examined neuronal responses in visual areas V1 and LM. Neurons often showed a preference for the left or right eye where a monocular grating was presented, and their responses were modulated by presenting a conflicting flashing Mondrian stimulus to the other eye. The direction of this modulation depended on whether neurons preferentially responded to contralateral or ipsilateral gratings. Responses of cells with a preference for ipsilateral gratings were suppressed during binocular conflict, while contralateral-preferring cells were less suppressed or even enhanced by binocular conflict. Binocular conflict modulation continued to occur under anesthesia, but response recovery after intermittent binocular conflict only occurred in awake mice. Responses to Mondrians were modulated by binocular conflict in a similar fashion as those to grating stimuli, which suggests that mouse binocular conflict processing exhibits a distinctive dependency on stimulus laterality over other stimulus features. Finally, a canonical binocular rivalry model could be successfully fitted to our data but lacked sufficient competitive inhibition and adaptation strength to create oscillatory activity under binocular conflict.

## Introduction

Despite decades of neuroscience research, the neural basis of conscious visual perception is still unclear (Koch et al., 2016; Pennartz, 2018; Seth & Bayne, 2022). Binocular rivalry (BR) has been widely used to study conscious perception in humans and non-human primates (Blake et al., 2014; Maier et al., 2012). BR occurs when incongruent images are presented to the left and right eye, which may cause a viewer’s perception to spontaneously alternate between the two images instead of fusing them (Alais & Blake, 2005). Previous studies in humans (Wheatstone, 1838) and macaques have shown that this phenomenon occurs in both species (Myerson et al., 1981) but whether it occurs in rodents is unknown. Although rodents have a smaller binocular field of view compared to primates (Priebe & McGee, 2014), they preferably process important stimuli through their binocular field, suggesting that binocular vision is important for rodent perception (Holmgren et al., 2021; Wallace et al., 2013). Because a wide array of brain recording and genetic manipulation techniques is available for use in rodents, rodents could serve as a model organism to study binocular conflict phenomena (Storm et al., 2017).

Here, we investigated whether mouse neuronal processing of binocularly conflicting stimuli in early visual areas is similar to that of primates. This could indicate that they share neuronal mechanisms for binocular conflict resolution. In macaques, a subset of neurons in primary visual cortex (V1) shows suppression of their responses to their preferred stimulus if that stimulus is reported to be perceptually suppressed during BR (Bahmani et al., 2014; Keliris et al., 2010; Leopold & Logothetis, 1996). As a similar suppression occurs in comparable stimulation conditions under anesthesia (Xu et al., 2016), it has been suggested that binocular conflict processing in early primate visual areas may serve as an important preconscious basis for further processing in higher areas (Bahmani et al., 2014). Therefore, we examined whether mouse early visual areas exhibit similar binocular response modulations as those of primates, and whether these modulations are sensitive to loss of consciousness during anesthesia.

We investigated neural mechanisms of binocular conflict processing in mice using an adaptation of a binocular stimulation paradigm called Continuous Flash Suppression (CFS) (Tsuchiya & Koch, 2004, 2005). This paradigm employs a flashing Mondrian stimulus on one eye (the mask) in a binocular display and has been shown to be particularly effective at suppressing the stimulus on the other eye (the target) from awareness in primates (Tsuchiya et al., 2006). We exposed passively observing and anesthetized mice to binocularly conflicting stimuli consisting of an achromatic flashing Mondrian mask and a drifting grating target, while we imaged calcium activity in cortical layer 2/3 cells in V1 and the lateromedial area (LM). We find that binocular conflict can suppress neuronal target responses in mouse V1 and LM to comparable degrees as in primate V1. While this suppression also occurred under anesthesia, recovery to pre-conflict levels of activity after binocular conflict offset was only apparent during wakefulness. Unlike primate V1, we find a strong dependency of the directionality of the response modulation on the eye exposed to the preferred grating stimulus. Independently of whether the preferred stimulus consisted of a target or a mask, cells with their preferred stimulus on the contralateral eye tended to only weakly suppress or even enhance their responses during binocular conflict, while responses of cells with a preference for ipsilateral stimuli were more strongly suppressed. Thus, mouse binocular conflict processing exhibits a distinctive dependency on stimulus laterality over other stimulus features. Finally, our flash suppression data could be fitted by a computational BR model, but given the best fitting parameter values, the model did not generate sufficient competitive inhibition and adaptation strength for oscillatory activity during binocular conflict in mouse visual cortex.

## Results

To study neuronal processing of binocular conflict in mice, we developed an apparatus that presents different images to the left and right eye of a mouse (Figure 1A; for similar designs see (Fu et al., 2023; Samonds et al., 2019)). While presenting two independent monocular images, we recorded the activity of cortical layer 2/3 neurons of mice using a two-photon microscope and recorded both eyes using cameras (Figure 1B, C). Based on widefield retinotopic mapping, we targeted the binocular region at the border of V1 and neighboring area LM for two-photon imaging (Figure 1C - F).

**Figure 1.**
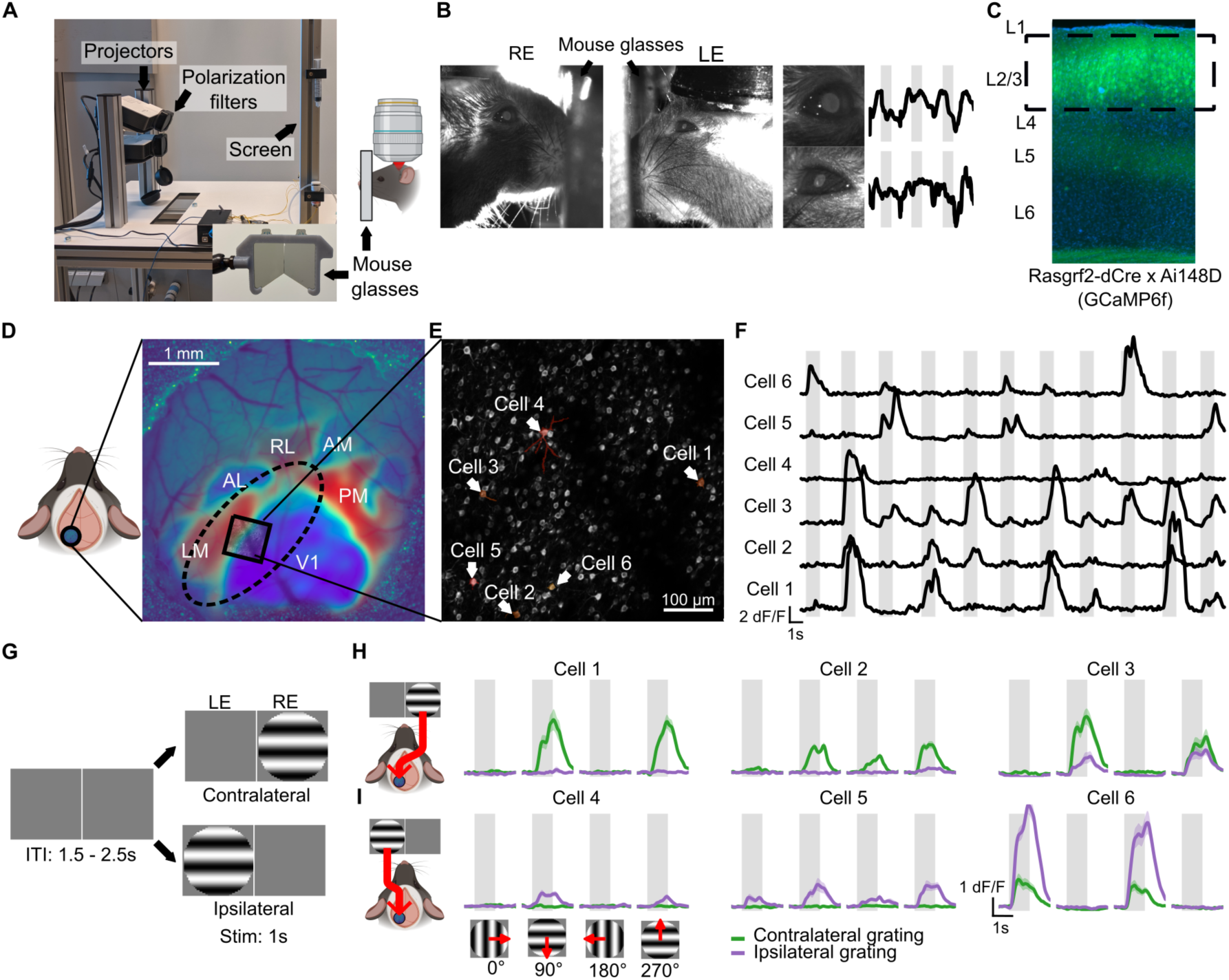
Binocular stimulation of mouse eyes and in vivo two-photon recordings. (A) Experimental setup to present binocularly conflicting stimuli to mice. The mouse drawn on the right is positioned adjacent to the screen (photograph on left). (B) Example frames from binocular eye tracking. (left) Position of mouse glasses. (right) Pupil dilation of each eye (a.u.) over 10s with stimulus periods denoted by grey shaded regions. RE: right eye, LE: left eye. (C) Fluorescence image of DAPI-stained cortical section showing expression of GCaMP6f in layer 2/3. Green = GCaMP, Blue = DAPI. (D) Schematic of cranial window position (left hemisphere) and widefield image (dark blue to yellow) overlaid with retinotopic map (blue, red) and an example imaging site (black square). The colors of the retinotopic map indicate the visual field sign at each pixel. Patches of shared field sign values delineate the location of a visual area (see also Methods). Dashed ellipse indicates the approximate outline of the lateral binocular zone. (E) Maximum-projection image (greyscale, each pixel depicts the highest fluorescence value of the recording) of example imaging session with six highlighted cells (orange to red). (F) Baseline-scaled (dF/F) fluorescence traces of the six cells highlighted in (E). Shaded areas represent periods of stimulus presentation. (G) Stimulation paradigm to assess selectivity for monocular gratings. Monocular gratings were shown with four different orientations (0, 90, 180 and 270**°**) to either the left or the right eye. Only two out of eight monocular gratings are depicted. ITI: intertrial interval. (H) Average responses of three example cells from E, F with a preference for gratings on the contralateral eye. Shaded region around curves represents SEM. Shaded grey areas represent stimulus presentation periods. (I) As (H) but now example cells with a preference for gratings on the ipsilateral eye.

### Binocular conflict suppresses target responses in mouse early visual areas

In a first experiment, we presented mice with short (1s) monocular or binocular stimuli. Each neuron’s orientation and eye selectivity were determined using eight monocular drifting gratings presented for 1s (four orientations x two eyes; Figure 1G - I). We then identified a neuron’s preferred monocular grating as the grating that elicited a neuron’s maximum average response. All gratings, regardless of the stimulated eye, were presented at the same central location in the visual field with a size of 30 visual degrees.

Inspired by (Tsuchiya et al., 2006), we investigated whether responses of neurons to their preferred monocular grating (the target) could be modulated by presenting a flashing, achromatic Mondrian stimulus (the mask) on the non-preferred eye at the same location in the visual field as the grating (Figure 2A). In primates, a small fraction of neurons (∼20%) in early visual areas suppresses their responses to their preferred stimulus during BR when the preferred stimulus is reported to be perceptually suppressed (Leopold & Logothetis, 1996) and such neuronal suppression also occurs when the non-preferred stimulus is flashed in a binocular display while the preferred stimulus is held constant (Bahmani et al., 2014; Keliris et al., 2010). Thus, we hypothesized that CFS-inspired binocular conflict could evoke neuronal target response suppression in early mouse visual areas (Figure 2B).

**Figure 2.**
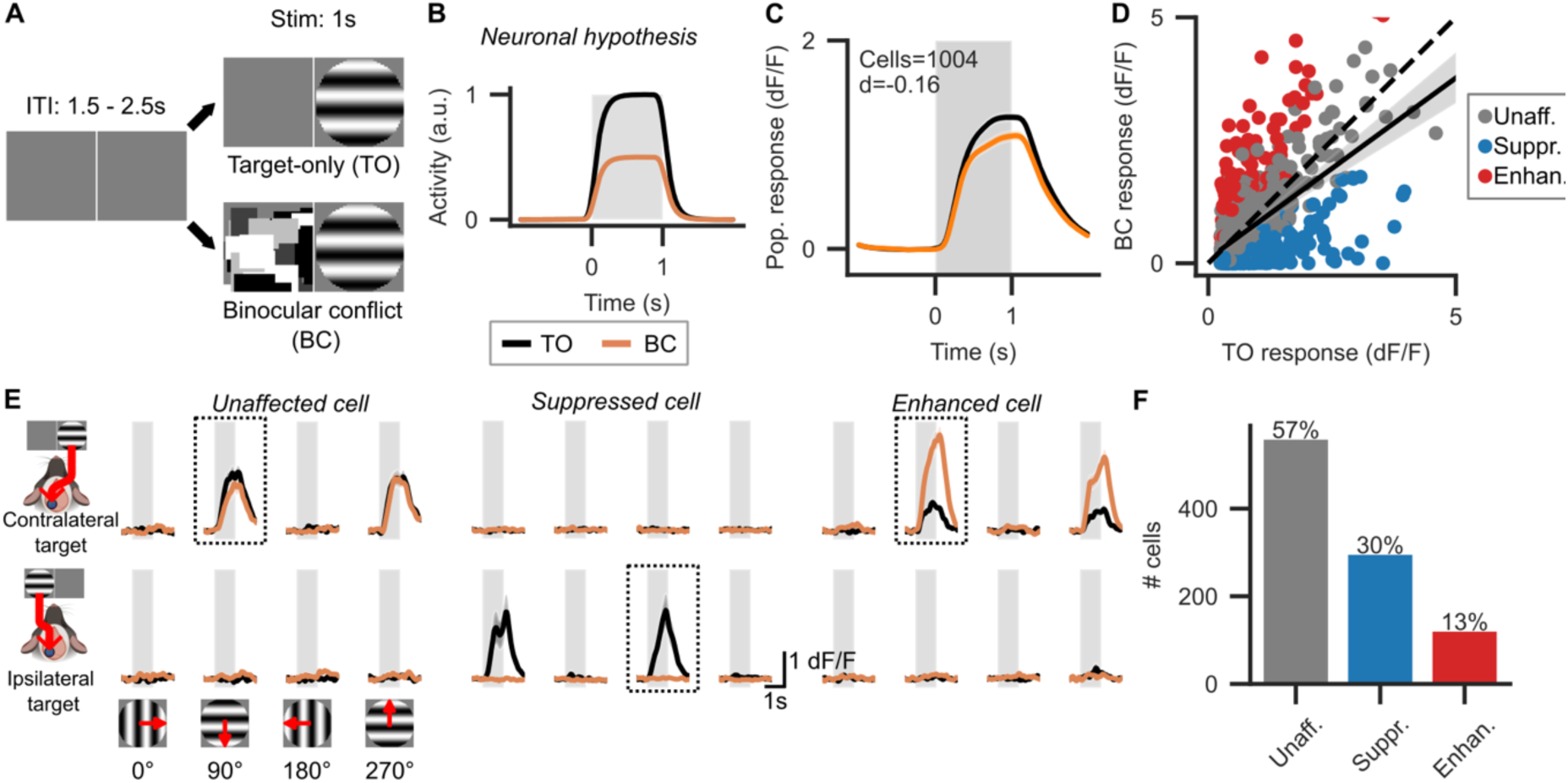
Flashing binocular conflict weakly suppresses the population response of target-selective cells in mouse V1 and LM during wakefulness. (A) Simultaneous-onset stimulation paradigm displaying binocularly conflicting stimuli. (B) Neuronal hypothesis: responses to preferred gratings (“targets”) are suppressed when in binocular conflict with a flashing achromatic Mondrian (“mask”). Shaded grey area indicates time of stimulus display. (C) Population response of V1 and LM cells to target-only (TO) and binocular conflict (BC) conditions. Shaded areas around lines indicate SEM (barely visible). Shaded grey rectangle indicates stimulus presentation period. d: Effective difference between TO and BC responses as quantified by Cohen’s d. (D) Scatterplot of average responses of individual cells to TO and BC condition. Each dot represents one cell. Dashed line: same response in both conditions. Solid line: linear regression, shaded area represents 95% confidence intervals from bootstrapping. (E) Average responses of three example cells to all eight TO and corresponding BC conditions. Plotting conventions as in C. Dotted rectangles indicate conditions with preferred target. (F) Bar plot indicating absolute number of V1 and LM cells with either no modulation during binocular conflict, significant suppression or significant facilitation. Fractions of cell counts are given as percentages on top of the bars.

We recorded neuronal activity at the V1-LM border of eight passively observing mice in 45 sessions. Stimuli either consisted of a monocular drifting grating (target-only condition, TO), a flashing monocular Mondrian (mask-only condition, MO) or both presented in binocular conflict (binocular conflict condition, BC). We restricted our analysis to cells that selectively responded to the targets and responded strongly to their preferred target (n = 1004 target-selective cells, see Methods). For each cell, we only analyzed TO and BC trials in which the target consisted of the cell’s preferred monocular grating. When comparing the population TO response with the BC response, we found the BC response to be weakly but significantly suppressed (Cohen’s d = −0.16, p < 0.001, two-sided Wilcoxon signed-rank test [tsWSRT], n = 1004 cells; Figure 2C). We next examined response modulations on the level of individual cells. We compared the average responses of each cell in TO trials with those in BC trials (two-sided Mann Whitney U test [tsMWUT], n = 40 trials, FDR correction for m = 1004 cells, α = 0.05) and found a large heterogeneity in significant response modulations with a bias towards suppression (Figure 2D – E). Specifically, while most cells’ responses were unaffected by the addition of the mask (570 / 1004 cells, 57%), 306 cells (30%) were suppressed, and a smaller fraction (128 cells, 13%) was enhanced (Figure 2F).

### Directionality of binocular response modulation depends on laterality of preferred target

Although the effect of flashing binocular conflict on target responses was net suppressive, we wondered why it was suppressive on some cells but facilitatory on others. To test whether this effect was dependent on stimulus laterality, we grouped cells by ocular dominance, that is, whether their preferred target was on the contra- or ipsilateral eye. We refer to these cells as ipsi- and contra-preferring cells in the following. In accordance with earlier studies reporting dominance of contralateral signals in mouse binocular visual cortex (Dräger, 1975; Salinas et al., 2017), we found more contra- (731 cells, 73%) than ipsi-preferring cells (273 cells, 27%). At first sight, ipsi-preferring cells tended to be more suppressed than contra-preferring cells (Figure 3A). Further analysis showed that this was true both on the level of population responses and individual cells. While the population response of contra-preferring cells was barely suppressed (Cohen’s d=-0.06, p < 0.001, tsWSRT, n = 731 cells), we found moderate to strong suppression of the ipsi-preferring population (Cohen’s d=-0.48, p < 0.001, n = 273 cells; Figure 3B-D). For individual cells, we found that about half of all ipsi-preferring cells were suppressed (142 cells, 52%) while only 22% of the contra-preferring cells (164 cells) showed suppression (Figure 3E-F). Finally, while target orientation decoding performance was worse during BC trials than TO trials for targets on either eye (Figure 3G), decoding performance during BC trials decreased more strongly for ipsi-than contralateral targets (difference in decoding accuracy between TO and BC trials for target on contralateral eye : −0.09 ± 0.01 [mean ± sem], ipsilateral eye: −0.16 ± 0.02, p < 0.001, tsWSRT).

**Figure 3.**
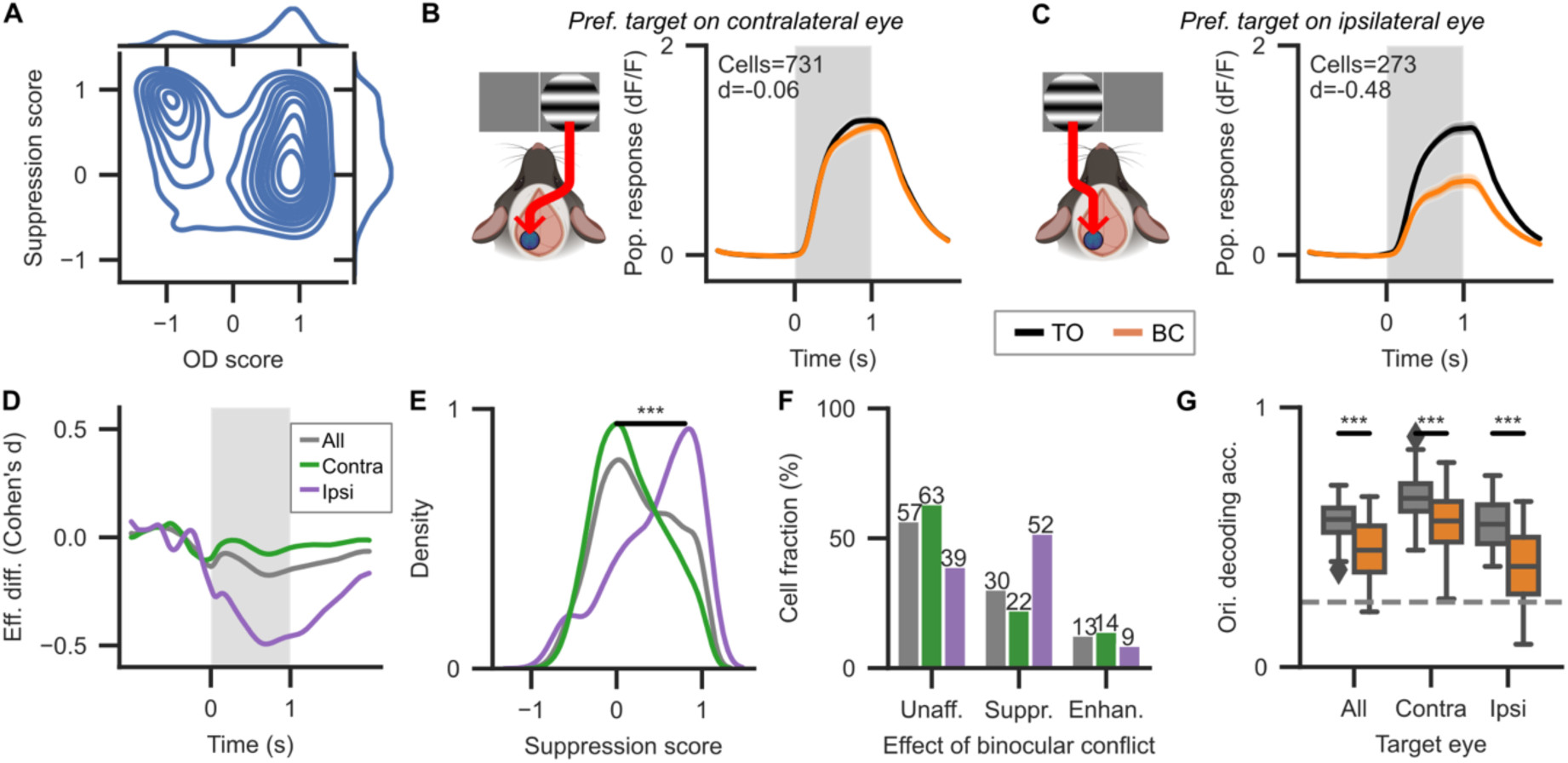
Cells preferring ipsilateral targets are more strongly suppressed by binocular conflict than cells preferring contralateral targets. (A) 2D Kernel density plot of ocular dominance (OD) scores and suppression scores (see Methods). An OD score of 1 indicates that a cell responds to a target of its preferred orientation only on the contralateral (right) but not the ipsilateral (left) eye. A suppression score of 1 indicates that a cell responds to its preferred target alone (TO condition) but does not respond at all when its preferred target is in binocular conflict with a flashing Mondrian (BC condition). (B) Population response of cells with their preferred target on the contralateral eye. Plotting conventions: see Figure 2C. d: Effective difference between TO and BC responses (Cohen’s d). (C) Same as B but for cells with their preferred target on the ipsilateral eye. (D) Effective difference between TO and BO population responses separated according to the eye of origin of their preferred target. All: any eye of origin, meaning all cells, Contra: cells with preferred target on contralateral eye, Ipsi: cells with preferred target on ipsilateral eye. (E) Kernel density plot of suppression scores according to the eye of origin of a cell’s preferred target. *** = p < 0.001, tsMWUT between suppression scores of contra- and ipsi-preferring cells. Colors as in D. (F) Fractions of significantly modulated cells. Numbers on top of bars indicate percentages. Colors as in D. (G) Target orientation decoding accuracy for TO (grey) and BC (orange) trials split by target eye. Target orientation was decoded for each recording session separately (see Methods). Targets were shown in four orientations. Dashed horizontal line indicates chance level (25%). Colors as in D. TO decoding performance exceeded BC decoding performance for all splits. *** = p < 0.001, tsWSRT, n = 45 recordings. Ori. decoding acc. = balanced orientation decoding accuracy.

We checked whether eye movements could explain this laterality effect but found little systematic eye movement to our stimuli (Supplementary figure 1). Furthermore, we were unable to distinguish V1 from LM cells based on their response patterns to all stimulus conditions (Supplementary figure 2). Therefore, we decided to pool neurons from both areas for the remainder of this study.

### Response suppression during binocular conflict persists under anesthesia

While response modulations during BR in early primate visual areas were originally directly linked to conscious perception (Leopold & Logothetis, 1996), recent studies have shown that they persist under anesthesia (Bahmani et al., 2014; Xu et al., 2016). To investigate whether response modulations in mouse visual areas depended on being in conscious, we repeated the above experiment while mice were under isoflurane-xylazine anesthesia. We recorded data from five mice (a subset of the previous 8 mice) in 11 sessions and identified 127 target-selective cells. This time, we found roughly equal fractions of contra- and ipsi-preferring cells (contra: 61 cells, 48%; ipsi: 66 cells, 52%). The BC population response under anesthesia was strongly suppressed compared to the TO response (Cohen’s d = −0.97, p < 0.001, tsWSRT, n = 127 cells; Figure 4A). Furthermore, response suppression occurred both in the contra- and ipsi-preferring population responses under anesthesia (both p < 0.001, n_contra_ = 61 cells, n_ipsi_ = 66 cells). Still, the ipsi-preferring population response was more strongly suppressed than its contra-preferring counterpart (Cohen’s d_contra_ = −0.69, d_ipsi_ = −1.22; Figure 4B–D) and suppression scores were significantly higher for ipsi- than contra-preferring cells (contra: 0.35 ± 0.04 [mean ± sem], ipsi: 0.55 ± 0.04, p < 0.001, tsMWUT).

**Figure 4.**
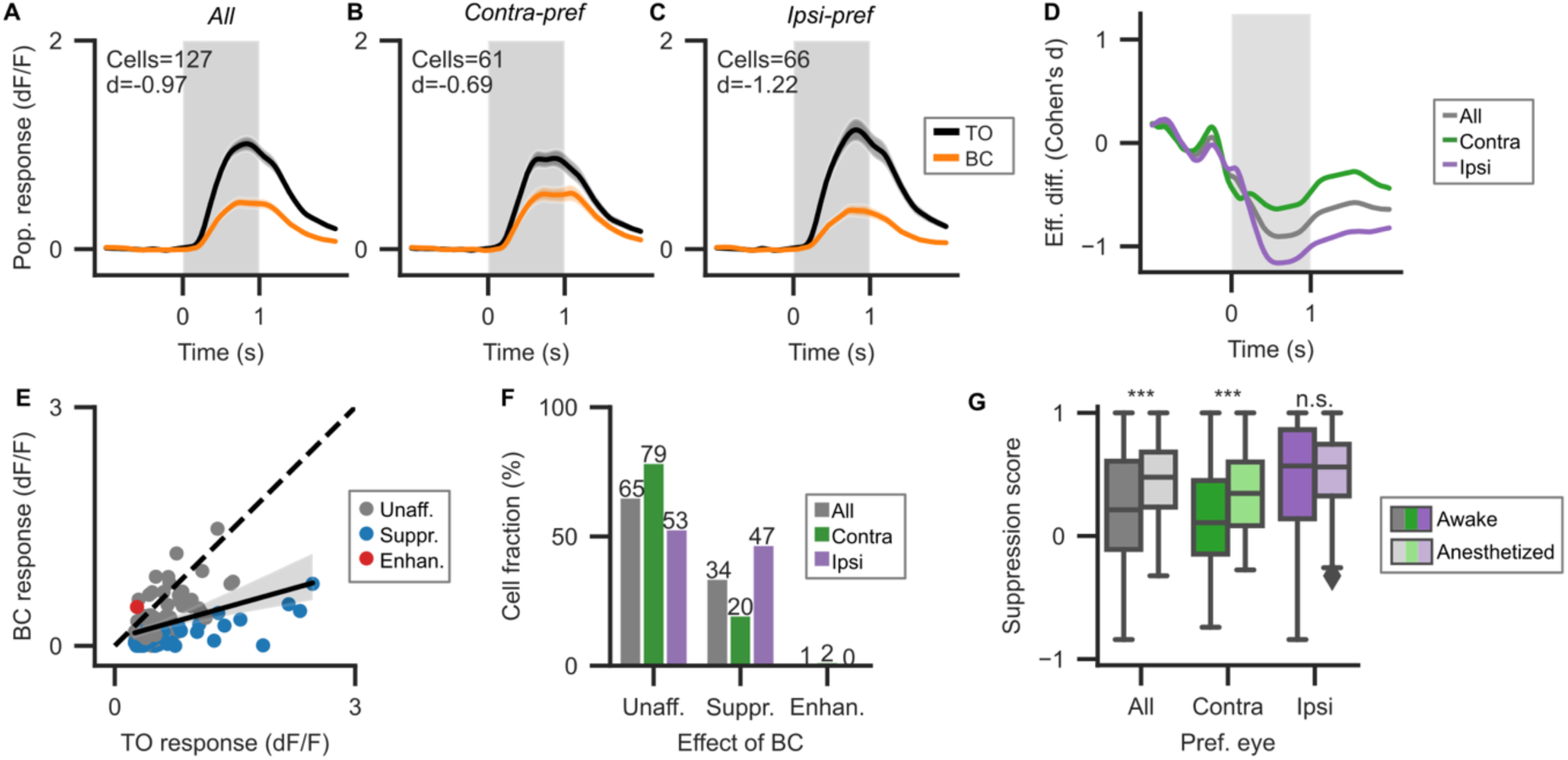
Binocular conflict responses are strongly suppressed under isoflurane-xylazine anesthesia. (A) Average population response to target-only (TO) and binocular conflict (BC) conditions under anesthesia. Plotting conventions as in previous figures. (B) Same as A but only for cells with their preferred target on the contralateral eye. (C) Same as A but only for cells with their preferred target on the ipsilateral eye. (D) Effective difference (Cohen’s d) between TO and BC population responses under anesthesia. Plotting conventions as in Figure 3D. (E) Scatterplot of average responses of individual cells under anesthesia. Plotting conventions as in Figure 2D. (F) Bar chart of cell fractions without and with significant modulation under anesthesia. Plotting conventions as in Figure 3F. (G) Boxplot of suppression scores in awake and anesthetized conditions. *** = p < 0.001, tsMWUT between awake and anesthetized suppression scores. n.s.: not significant, p > 0.05. For suppression scores, see Fig. 3D.

When looking at response modulations of individual cells, we found similar fractions of suppressed cells in contra- and ipsi-preferring populations as during wakefulness (contra: 12 cells, 20%, ipsi: 31 cells, 47%). Unlike during wakefulness, we found almost no cells with response enhancement under anesthesia (1 cell = 1%; Figure 4E-F). When comparing wakefulness to anesthesia, there was no statistically significant difference in the amount of suppression of the ipsi-preferring cells (p = 0.538, tsMWUT, n = 339 cells) but the amount of suppression in contra-preferring cells was significantly higher during anesthesia than wakefulness (p < 0.001, n = 792 cells; Figure 4G).

### Neuronal responses recover to undisturbed level after binocular conflict offset in awake mice

After observing that neuronal target responses are modulated when mask and target are presented simultaneously, we investigated what happens to such responses when the mask is intermittently shown during a constant target display. This delayed-onset paradigm can be considered a combination of binocular flash suppression (Wolfe, 1984) and CFS (Tsuchiya et al., 2006). In a new experiment, we presented mice with two stimulus conditions: (1) a monophasic “long target-only” condition (LTO) in which a monocular drifting grating target (either 0 or 90° orientation) was shown for 6s and (2) a triphasic “intermittent binocular conflict” condition (IBC) in which a target was shown for 6s on one eye while a flashing achromatic Mondrian mask appeared 2s after target onset for 2s on the other eye (Figure 5A). In primates, neuronal response suppression during BR is thought to cease when a subject becomes aware of the previously perceptually suppressed preferred stimulus of those neurons (Tong et al., 2006). We therefore investigated whether neuronal responses in mouse V1-LM during IBC trials would recover back to the level of LTO trials after binocular conflict offset (Figure 5B). We collected data from four mice (a subset of the 8 mice of the first experiment) in 16 sessions and identified 510 target-selective cells in V1 and LM. Three example cells in this delayed-onset paradigm are shown in Figure 5C.

**Figure 5.**
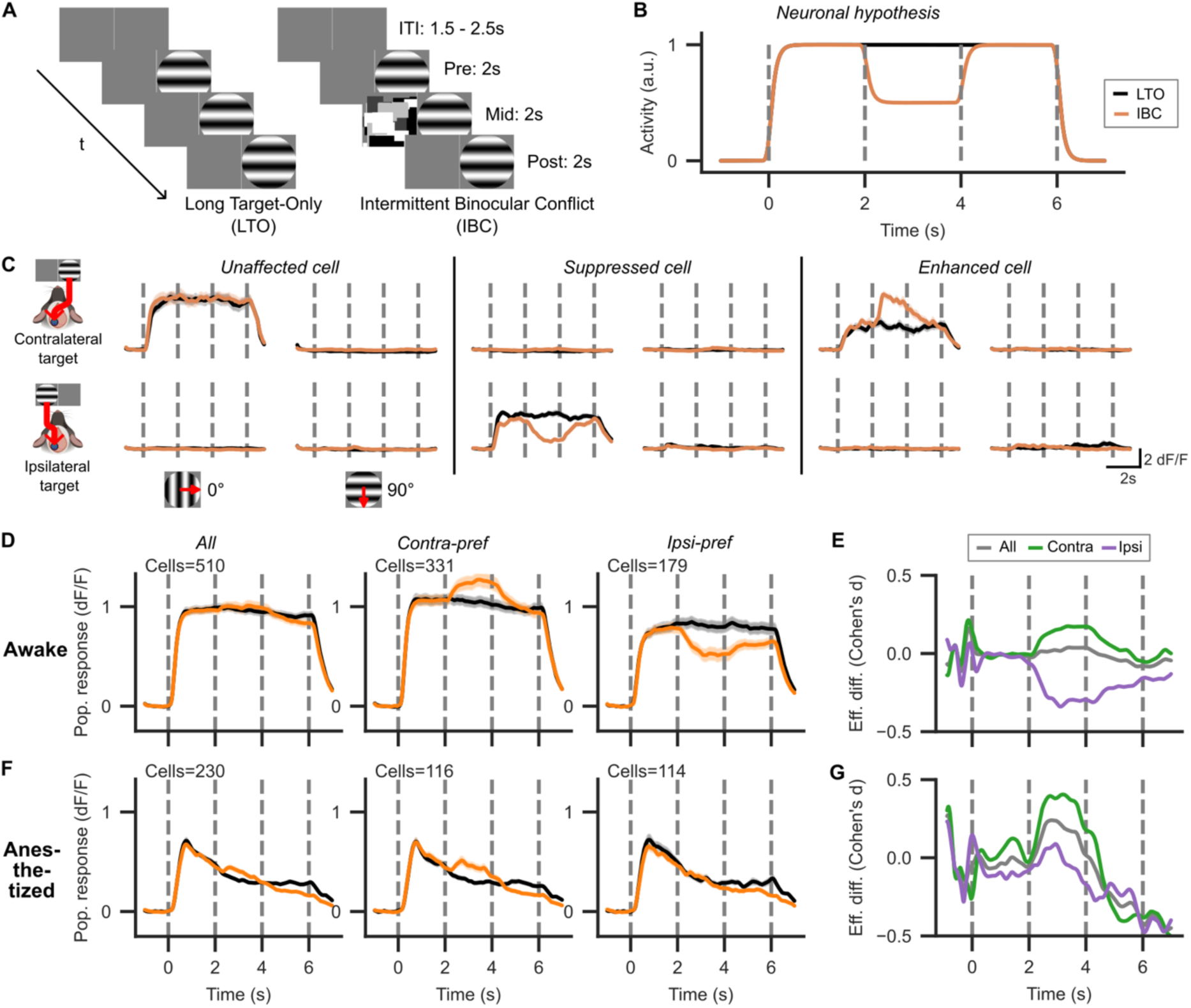
Population responses recover after binocular conflict offset during wakefulness but not anesthesia. (A) Delayed-onset stimulus paradigm with intermittent binocular conflict. (B) Neuronal hypothesis: during wakefulness, we expected neuronal responses to be suppressed during binocular conflict but to return to the level of monocular responses after binocular conflict offset. (C) Average responses of three example cells to LTO and IBC conditions. (D) Population responses to LTO and IBC conditions during wakefulness. Plotting conventions as in previous figures. Dashed lines indicate onset of a new stimulus phase. (E) Effective difference (Cohen’s d) between LTO and IBC population responses during wakefulness. (F) Same as D but now under anesthesia. (G) Same as E but now under anesthesia.

Overall, we found no significant difference when comparing LTO and IBC population responses during binocular conflict (Cohen’s d = +0.03, p = 0.064, tsWSRT, n = 510 cells; Figure 5D, left). When dividing the overall population into contra- and ipsi-preferring cells, we found opposing modulations: the contra-preferring population response was enhanced during binocular conflict (Cohen’s d = +0.17, p < 0.001, tsWSRT, n = 331 cells) while the ipsi-preferring population response was suppressed (Cohen’s d = −0.31, p < 0.001, n = 179 cells; Figure 5D, middle & right). After offset of the mask but not the target in IBC trials (so after t = 4s), both contra- and ipsi-preferring cells converged back to the level of LTO trials (Figure 5D-E). Concretely, the contra-preferring IBC response fully converged to the LTO level (Cohen’s d = −0.001, p = 0.318, tsWSRT). The ipsi-preferring IBC response was still suppressed (Cohen’s d = −0.20, p < 0.001, tsWSRT) but shifted significantly towards the LTO level (suppression scores during conflict: 0.35 ± 0.03 [mean ± sem], after conflict: 0.17 ± 0.03, p < 0.001, tsWSRT).

### Neuronal responses after binocular conflict offset fail to recover under anesthesia

We repeated this experiment under isoflurane-xylazine anesthesia to see whether response recovery following binocular conflict offset was dependent on conscious state. From the same four mice, we recorded data in 13 anesthetized sessions and identified 230 target-selective cells in V1 and LM. Again, we found no significant differences between LTO and IBC responses during binocular conflict in the overall population response (Cohen’s d = +0.16, p = 0.103, tsWSRT, n = 230 cells). Contrary to the first anesthetized experiment we found that response enhancement of the contra-preferring population during binocular conflict persisted under anesthesia (Cohen’s d = +0.35, p < 0.001, tsWSRT, n = 116 cells) while response suppression of the ipsi-preferring population was attenuated (Cohen’s d = −0.06, p = 0.254, n = 114 cells; Figure 5F). Strikingly, neither population’s average response converged back to the level of LTO trials after mask offset, with IBC responses of both populations falling below the level of corresponding LTO responses (Figure 5F-G; contra-preferring population: Cohen’s d = −0.40, ipsi-preferring population: Cohen’s d = −0.28, both p < 0.001, tsWSRT).

We checked for eye movements that could bias these results but again found few, unsystematic eye movements to our stimuli (Supplementary Figure 3). An additional control condition in which, instead of presenting intermittent BC, the target stimulus was intermittently removed from display 2s after target onset for 2s showed that the level of suppression reached during intermittent BC was not as strong as the drop in population response occurring when the target was temporarily removed from the display (Supplementary Figure 4).

### Mask responses are also suppressed by binocular conflict

In humans, perceptual dominance durations during BR have been linked to stimulus saliency (Levelt, 1965). For example, it has been shown that a flashing Mondrian can suppress a Gabor patch target from awareness for prolonged periods in humans during CFS but not vice versa (Tsuchiya et al., 2006). Although it is unclear whether our flashing Mondrian masks would be more salient to a mouse than the drifting grating target, as both are rather abstract stimuli, we investigated whether, besides stimulus laterality, response modulations during binocular conflict in mice were dependent on stimulus type (Figure 6A). Specifically, we examined whether responses to ipsilateral masks would also be suppressed during binocular conflict with a target despite the masks having higher saliency by human standards (Figure 6B). For this analysis, we re-examined the data from the first experiment (Figure 2, 3) during wakefulness but now selected cells with a preference for one of the two masks (see Methods). We identified 714 mask-selective cells (Figure 6C). Just as neurons with a preference for ipsilateral targets suppressed their responses when adding a contralateral mask, neurons with a preference for ipsilateral masks suppressed their responses when adding a target (Cohen’s d=-0.54, p < 0.001, tsWSRT, n = 210 cells; Figure 6E). The suppression in the ipsi-preferring population of Mondrian-selective cells was again greater than its contra-preferring counterpart (contra-pref. suppression score = 0.10 ± 0.01 [mean ± sem], ipsi-pref. suppression score = 0.32 ± 0.03, p < 0.001, tsMWUT). We therefore conclude that in our data, the directionality of response modulations is largely determined by stimulus laterality: across all stimulus types and conscious states, response modulations of contra-preferring cells were consistently comparatively more positive than those of ipsi-preferring cells.

**Figure 6.**
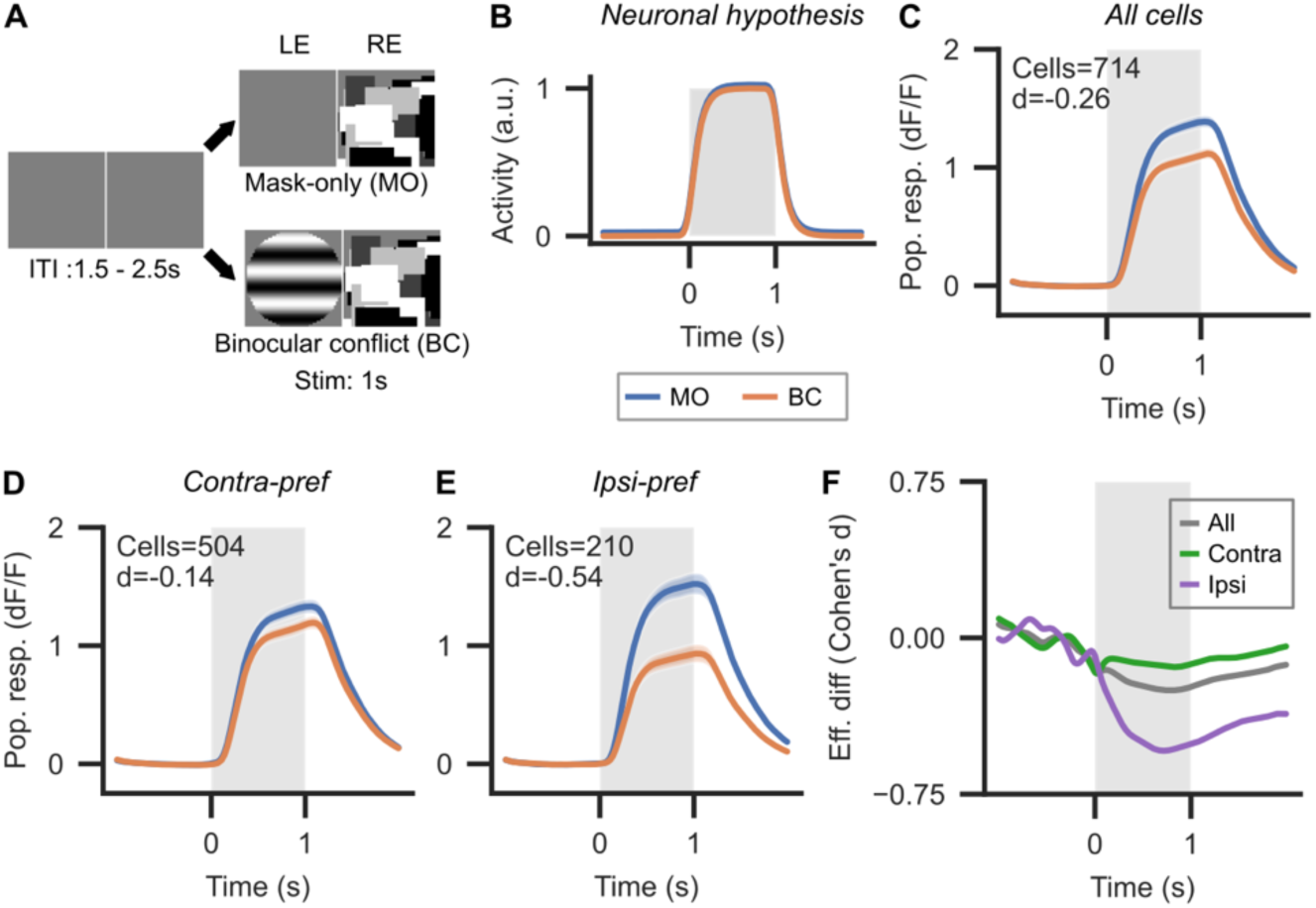
Ipsilateral response suppression during wakefulness is independent of stimulus type. (A) Stimulation paradigm to investigate response modulations of mask-selective cells (cf. Fig. 2). (B) Illustration of neuronal hypothesis. We expected mask responses to be unaffected by binocular conflict. (C) Population responses to mask-only (MO) and binocular conflict (BC) conditions. Plotting conventions as in previous figures. (D) Same as C but now for cells with preferred mask on contralateral side only. (E) Same as C but now for cells with preferred mask on ipsilateral side only. (F) Effective difference (Cohen’s d) between MO and BC population responses.

### An attractor network models binocular suppression but predicts insufficient inhibition for spontaneous binocular rivalry-like oscillations in mouse visual cortex

To understand whether our results are consistent with canonical attractor network models of BR in humans and non-human primates (NHPs), we fit a minimal model of BR developed by (Wilson, 2007) to the neuronal data of the simultaneous-onset experiment (Figure 7A). We chose the model of (Wilson, 2007) as it is arguably the simplest model of BR in the literature that includes the key canonical features common across BR models: mutual inhibition between monocular populations and slow hyperpolarizing spike-dependent adaptation (Shpiro et al., 2007). As the model was developed to account for the mesoscale mean-field firing rate behavior of large neuronal populations (Wilson & Cowan, 2021), we focused on the grand population average results without considering laterality differences.

**Figure 7.**
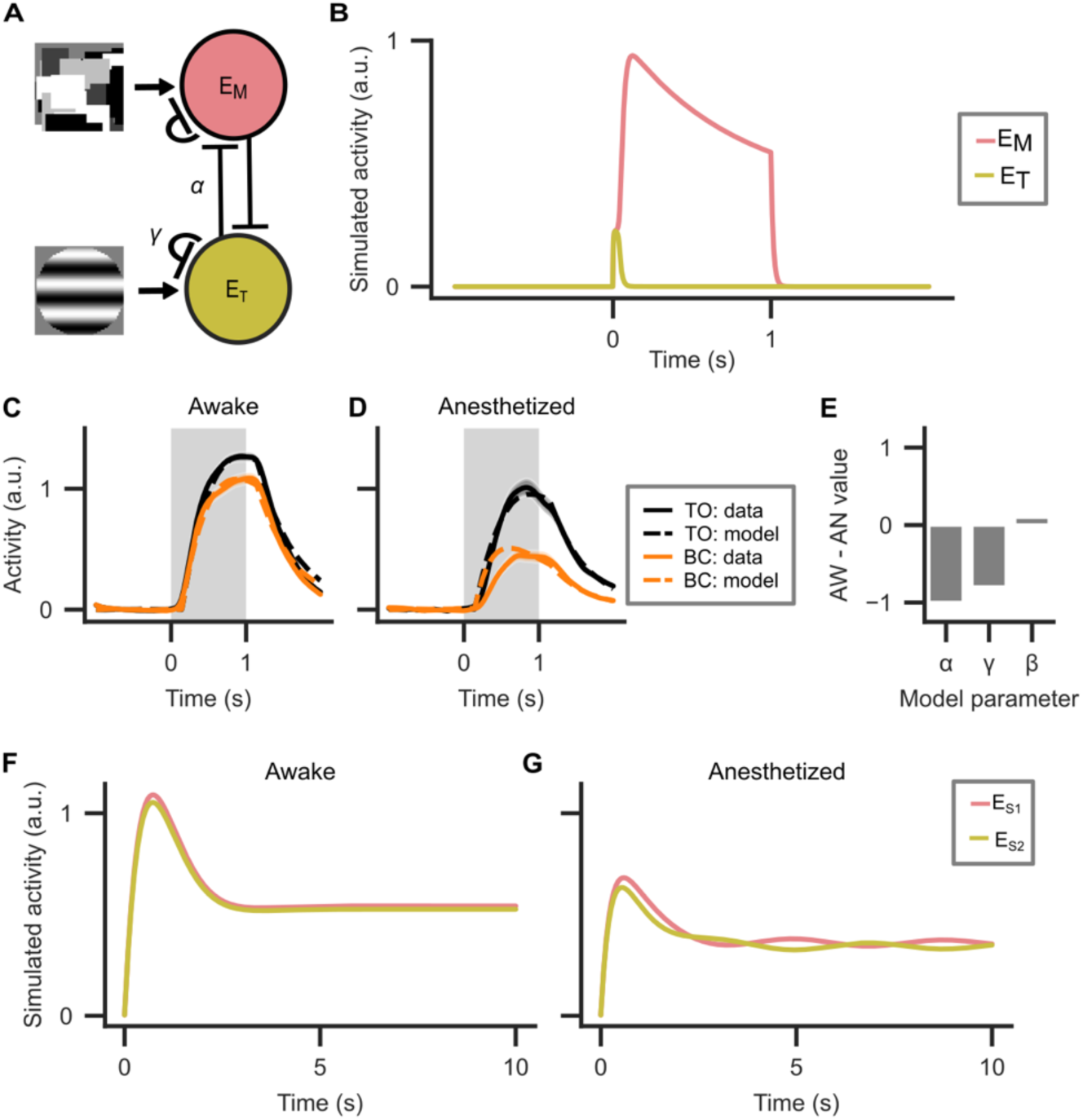
Attractor network model fits data on binocular conflict with simultaneous onset and predicts insufficient inhibition to generate oscillations. (A) Model architecture of BR attractor network model based on (Wilson, 2007). α = competitive inhibition parameter, γ = activity-dependent adaptation parameter, E_M_ = population activity of mask-driven population, E_T_ = population activity of target-driven population. Each population also has a transfer function (not shown in figure) governed by the gain parameter β. (B) Model dynamics during BC condition when using parameters from human CFS literature. The activity of the target-driven population is almost completely suppressed. (C) Mouse visual cortex data and model fit for BC condition during wakefulness. Grey shaded rectangle indicates stimulus presentation period. (D) Like (C) but during anesthesia. (E) Difference in model parameters between fits to awake (AW) and anesthetized (AN) data. Competitive inhibition (α) and spike-dependent adaptation (γ) parameters are higher during anesthesia than wakefulness. (F) Simulated activity of two stimulus-driven populations (E_S1_, E_S2_) during binocularly conflicting stimulation using best-fit parameters during wakefulness. Here, S1 and S2 are two equal-strength monocular stimuli that would elicit BR in humans and NHPs (e.g. two orthogonal gratings). No oscillations occur during wakefulness due to insufficient inhibition and hyperpolarizing adaptation current. (G) Like (F) but using best-fit parameters during anesthesia. Although inhibition and adaptation are increased, only minor oscillations appear in the simulated data.

The model had five free parameters representing (1) reciprocal inhibition between two competing populations, (2) spike-dependent adaptation, (3) activation function gain and (4, 5) the strength of mask and target stimuli. Using parameters from a previous modeling study of human CFS (Whyte et al., 2025) we observed total dominance of the simulated mask-driven population activity over the target-driven population activity during BC stimulation (Figure 7B). We then fit the same model to our own data using a multi-step grid search (see Methods). Parameters were fitted separately to awake and anesthetized data. After fitting, the attractor models reproduced our data well (Figure 7C-D). In the fit to the anesthetized data we saw an increase in the effects of competitive inhibition and adaptation, as well as a small reduction in the gain of the transfer function (Figure 7E). This is consistent with a known mechanism of action of isoflurane (as a GABA_A_ receptor agonist) (Banks & Pearce, 1999) and suggests that the increase in suppression under anesthesia may be the result of increased competitive inhibition and activity-dependent adaptation. Even with the increased competitive inhibition and adaptation strength, simulations of standard BR with the best-fitting parameters during both wakefulness and anesthesia failed to show rivalry-like oscillatory dynamics (Figure 7F-G). Thus, although binocular conflict modulations do occur in the mouse, the strength of competitive inhibition and hyperpolarizing adaptation may be insufficient to initiate the oscillatory activity that is thought to underlie BR in primates.

## Discussion

We presented binocularly conflicting CFS stimuli to awake and anesthetized mice and examined neuronal responses in V1 and LM. Responses of neurons to their preferred monocular grating target were often modulated by presenting a conflicting flashing Mondrian mask to the other eye. The direction of modulation depended largely on whether neurons preferentially responded to contralateral or ipsilateral targets. Cells with a preference for ipsilateral targets were suppressed during binocular conflict, while contra-preferring cells were less suppressed or even enhanced by binocular conflict. Response modulations by binocular conflict continued to occur under anesthesia, but response recovery after intermittent binocular conflict only occurred in awake mice. Binocular conflict modulated responses to masks in a similar fashion as those to grating stimuli, suggesting that binocular response modulations depend on laterality, but not stimulus type. Finally, a canonical BR model fitted our simultaneous-onset binocular conflict data well, but also predicted there is insufficient competitive inhibition and adaptation strength in mouse visual cortex to create BR-like oscillatory activity.

### Stimulus laterality in mouse visual cortex

While we were expecting to find binocular response modulations in mice based on previous studies in both mice and primates (Dougherty et al., 2019; Fu et al., 2023; Leopold & Logothetis, 1996; Scholl et al., 2013), we were surprised to find a dependency of the modulations on stimulus laterality. To our knowledge, similar findings have not been reported in comparable primate BR studies. One reason for this difference may lie in the projection pattern of retinal ganglion cells (RGCs). In primates, roughly half of all RGC axons cross over at the optic chiasm, leading to roughly equal amounts of contra- and ipsilateral signals in binocular V1 (Leinonen & Tanila, 2018). In mice, >95% of axons cross over at the optic chiasm such that even the binocular part of mouse visual cortex is dominated by contralateral signals (Dräger, 1975; Leinonen & Tanila, 2018). Because of this contralateral dominance, contralateral signals could be more robust to disturbances by binocular conflict than ipsilateral ones.

Another key difference between primates and mice is the notable absence of ocular dominance columns in mice (but see Goltstein et al., 2025). BR in primates may rely on competition between ocular dominance columns (Blake, 1989). It is tempting to speculate that ipsilateral response suppression could form the basis of an alternative mechanism of binocular conflict resolution in mammalian species without ocular dominance columns. Regarding such mechanisms, we however note two caveats: first, ipsilateral signals were only partially suppressed in our data (ipsilateral target orientation could still be decoded above chance in BC trials, Figure 3G). Second, because all recordings were performed in the left hemisphere, it would need to be verified that ipsilateral signals are also suppressed in the right hemisphere.

### Differences between simultaneous- and delayed-onset experiments

We found differences in response modulations by binocular conflict depending on whether target and mask were presented simultaneously or whether the mask was added with a delay. In awake mice during simultaneous-onset binocular conflict, the effect of binocular conflict was net suppressive while there was no net effect during delayed-onset binocular conflict. We also found opposing effects of anesthesia on response modulations depending on onset timing. During simultaneous onset, suppression was significantly higher during anesthesia than wakefulness (suppression score under wakefulness: 0.24 ± 0.01 [mean ± sem], anesthesia: 0.45 ± 0.03, Figure 4G). However, during delayed onset, suppression was attenuated during anesthesia (suppression score under wakefulness: 0.11 ± 0.02, anesthesia: −0.02 ± 0.03; data not shown). Many factors may play a role in explaining the differences between the two experiments including differences in cell and subjects counts. Next, we will discuss two factors mentioned in human and NHP BR studies.

First, the simultaneous-onset experiment investigates a phase of binocular conflict processing that in humans is referred to as ‘onset rivalry’ and is thought to differ significantly from prolonged rivalry. In humans, onset rivalry is characterized by a fusion of conflicting stimuli and perceptual biases for one over the other stimulus (Carter & Cavanagh, 2007; Wolfe, 1983). Both perceptual effects, even if they would translate to mice, are hard to relate to the neuronal findings. If delayed-onset experiments produced stronger perceptual suppression in mice, one would expect they also produced stronger neuronal suppression. If mice had a perceptual bias for one of the stimuli during simultaneous-onset experiments, one would expect modulations of target and mask to be markedly different (but they are not) or absent in the overall population because subjects’ biases cancel out (but they do occur). Second, primate BR has been linked to reciprocal suppression of the populations representing the two conflicting stimuli (Wilson, 2007) (Figure 7A) and delayed-onset BR experiments have been linked to adaptation. The target in our study was a drifting grating to which adaptation is probably limited (Goltstein et al. 2018). Reciprocal suppression could explain a suppression effect on both mask and target responses in the simultaneous-onset experiment where stimuli could be presented too briefly for one stimulus to achieve dominance over the other. Adaptation, on the other hand, does not explain our findings well: if we assume that because of adaptation, the suppressive effect of the target on potential rivaling stimuli has ceded, the mask would presumably achieve dominance more easily and suppress the target responses more strongly than during simultaneous-onset experiments – but this is not the case in our study.

### Attractor dynamics

With appropriately configured parameters, the attractor model was able to simulate our simultaneous-onset data under both awake and anesthetized conditions (Figure 7A). Isoflurane anesthesia is known to boost GABAergic inhibition and K+ current (Alkire et al., 2008; Franks, 2008) which is consistent with the increase in the adaptation and competitive inhibition parameters of the model, required to reproduce our simultaneous-onset observations under anesthesia (Figure 7E). The delayed-onset results appear to be more complicated than can be captured by the canonical model. To explain these results, we propose that, in addition to a reciprocal suppression mechanism, a population representing one of the two conflicting stimuli may consist of cells having recurrent excitatory connections. When comparing, for instance, the lack of suppression in the awake condition for all cells in the delayed-onset setup (Figure 5D) to the suppression of all cells in the simultaneous-onset setup (Figure 2C), the difference may be explained by an attractor-like stabilization of the grating representation in contra-preferring cells occurring before the Mondrian is displayed, consistent with the enhancement of grating responses in these cells during the conflict phase (Figure 5D, middle).

During anesthesia in the delayed-onset paradigm, population responses show an overall decline following the initial peak in the response to the first stimulus (Figure 5F), causing the suppression of the ipsi-preferring cells to be less apparent: there is less population grating response left to suppress, and the population representing the Mondrian pattern can be presumed to have become weaker at this stage as well. Strikingly, the response of the ipsi-preferring cells does not recover after the Mondrian has disappeared (Figure 5F, right panel), reinforcing the notion that reverberatory attractor properties have been weakened in that population. An involvement of recurrent excitatory connections is consistent with microcircuitry findings on mouse visual cortex (Ko et al., 2011) and with the antagonizing effect of isoflurane on of NMDA receptors (Alkire et al., 2008; Franks, 2008). Combined with our findings that response modulations were laterality-dependent, we conclude that a mechanism of reciprocal suppression between neuronal populations, each endowed with recurrent excitatory connections, offers a stronger potential for explaining our results than a mechanism chiefly relying on adaptation or inhibition. Future synaptically detailed modelling and experiments are needed to test this prediction.

### Comparison to macaque V1 recordings during BR

Electrophysiological studies of macaque V1 during BR or one of its variants, binocular flash suppression, have consistently found that a small fraction (20% - 25%) of neurons suppresses their responses to their preferred stimulus when that stimulus is perceptually suppressed (Bahmani et al., 2014; Keliris et al., 2010; Leopold & Logothetis, 1996). This response suppression is rather weak (Keliris et al., 2010: d’ = 0.41, Bahmani et al., 2014: d’ = 0.25) and much smaller than that elicited by physical removal of the preferred stimulus from the display. At least one study reported roughly similar fractions of suppressed and enhanced recording sites during BR in macaque V1 (Gail et al., 2004). Nonetheless, the effect of binocular flash suppression on macaque V1 population responses is net suppressive (Bahmani et al., 2014; Keliris et al., 2010). Finally, such suppression during binocular flash suppression also occurs in V1 under anesthesia (Bahmani et al., 2014).

In comparison, while we found a small net suppressive effect of flashing binocular conflict on target responses of mouse V1 and LM cells during simultaneous-onset binocular conflict, we did not observe such net suppression during delayed-onset binocular conflict (at least in the general population average). The latter finding is contrary to the small but robust suppression found in monkey V1 binocular flash suppression studies where one of the two stimuli was presented with a delayed onset. Nonetheless, we found roughly similar fractions of suppressed cells as reported in monkeys (30% during simultaneous onset, 15% during delayed onset). However, such fractions are sensitive to the type of multiple comparison correction used. Finally, while binocular modulations continued to occur under anesthesia in mouse V1 and LM, we found opposing effects of anesthesia on response modulations depending on whether the mask was presented simultaneously or with a delay. Taking the delayed-onset paradigm to be closest to macaque binocular flash suppression studies, our findings under anesthesia in the mouse run counter to those from macaque studies. As suggested by the model (Figure 7), the differences may be explained by a lower level of reciprocal inhibition in the mouse.

### Adaptation of Continuous Flash Suppression stimuli for mice

While adapting primate CFS stimulation paradigms to suit the features of mouse vision, we decided to use rather large stimuli (gratings: 30 visual degrees, Mondrians: 35 degrees). In humans, larger binocularly conflicting stimuli have been associated with an increase of episodes of piecemeal rivalry: fragmented perception of both the left-eye and right-eye stimuli together). Stimuli no larger than 10 visual degrees are therefore recommended in human BR studies (Carmel et al., 2010). Although it is unknown whether mice perceive binocular conflict in episodes of dominance-suppression or piecemeal rivalry, such larger stimuli could potentially have resulted in less clear neuronal response modulations. Nonetheless, there were good reasons to use comparatively large stimuli. While primates are foveates with high spatial acuity in their central visual field, mice are afoveates (but see van Beest et al., 2021) whose spatial acuity is about 60x lower than that of humans (Leinonen & Tanila, 2018). Furthermore, neurons in mouse V1 preferably respond to gratings with a much lower spatial frequency than the resolution limit allows (about 0.05 cpd, Marshel et al., 2011). We therefore settled on an intermediate spatial frequency (0.1 cpd) and stimulus size. While our stimuli lie well within the binocular field of the mouse (which starts at 35 - 40 visual degrees width in the horizontal plane and increases in width with elevation, Dräger, 1975; Samonds et al., 2019), it deserves further testing whether the fraction of modulated cells during binocular conflict correlates inversely with stimulus size.

### Comparison to two recent preprints on mouse binocular conflict

Two recent preprints have investigated binocular conflict processing in mouse V1, albeit with different stimuli than we used. The first (Montgomery et al., 2025) reports higher activity of mouse V1 L2/3 pyramidal cells when stimulated with incongruent rather than monocular gratings. We do not find a similar net facilitatory effect of binocular conflict but note the difference in incongruent stimulation (Montgomery et al.: incongruent gratings, this study: CFS) and in cell and condition selection in the analysis (Montgomery et al.: no checks for orientation tuning, averages include conditions without preferred grating, this study: orientation-tuned cells responding to conditions with preferred grating only; also LM cells). The second preprint (Timplalexi et al., 2025) reports evidence for spontaneous alternating activity of stimulus-tuned populations during stimulation with incongruent gratings in mouse V1 based on confidence measures of linear support vector machine decoders. This runs counter to the predictions of our model (Figure 7) and the findings that the contra-preferring population in our study remained relatively unaffected by binocular conflict. It is possible that Timplalexi et al.’s decoding method detects more subtle changes than revealed by population averages or that this oscillatory activity only emerges after longer incongruent stimulation.

### Outlook

Rodents have only recently become popular as model organisms in consciousness research (Storm et al., 2017). BR is an important tool to study visual awareness in primates (Blake et al., 2014) and if mice experience BR like primates (with a similar underlying mechanism), research on the neural basis of BR could be accelerated significantly by using mice as subjects. If mice do not experience primate-like BR when presented with binocularly conflicting stimuli, it does remain interesting to study how mice resolve binocular conflict perceptually. It will therefore be important to assess mouse perception of binocularly conflicting stimuli with behavioral testing in the future.

In primates, the fraction of cells with modulations during BR rises along the ventral visual processing stream (Hesse & Tsao, 2020; Leopold & Logothetis, 1996, 1999; Logothetis & Schall, 1989; Sheinberg & Logothetis, 1997). Future studies of neuronal response modulations could explore whether similar results hold in mice as well. Despite considerable differences between mouse and primate visual areas, recently, efforts have been made to establish a hierarchy of mouse visual areas with a similar division into ventral and dorsal streams (D’Souza et al., 2022). Laterointermediate (LI) and postrhinal (POR) areas lie at the top of the putative mouse ventral stream and could therefore be used to study binocular conflict. Another promising target is mouse prefrontal cortex because macaque prefrontal cortex has been shown to be strongly modulated during BR in both task-behaving and passive animals (Dwarakanath et al., 2023; Kapoor et al., 2022; Panagiotaropoulos et al., 2012).

## Acknowledgements

We thank Prof. Chris Schaffer and Daniel Rivera for providing hardware and software of the custom-made vitals monitor. We also thank Udo van Hes and Clint Janssen for their help with constructing the binocular setup.

## Methods

### Animals

All animal experiments were performed according to the national and institutional regulations. The experimental protocol was approved by the Dutch Commission for Animal Experiments (CCD application number: AVD11100202216078) and by the Animal Welfare Body of the University of Amsterdam. We used double transgenic mice (background: C57BL/6J) from in-house breeding of Rasgrf2-2A-dCre (Jax #022864, Song et al., 2017, Cre-driver line) and Ai148D (JAX #030328, Daigle et al., 2018, GCaMP6f reporter line) mouse lines aged between 8 and 52 weeks. We included data from n = 8 mice (3 females) in this study. Mice were housed socially on a reversed day night cycle (lights on: 8 pm, off: 8 am) and had free access to water and food. GCaMP6f expression was induced by injecting mice with 300 mg / kg trimethoprim (TMP T883-5G, Sigma-Aldrich) dissolved in dimethylsulfoxide (DMSO) intraperitoneally (IP, concentration: 250 mg/ml) once per day over a series of three days.

### Surgery

Mice were implanted with a custom-built titanium head bar for head fixation and a cranial window for imaging. For analgesic purposes, mice received a subcutaneous injection of 10 mg/kg carprofen (Carprofelican, Dechra) at the start of surgery. Anesthesia was induced using 4-5% isoflurane in 100% oxygen and isoflurane levels were lowered during surgery to 1%–2%. Anesthesia depth was confirmed by testing the pedal withdrawal reflex. The fur on the mouse head was shaved off and part of the skin above the mouse skull was surgically removed after treatment with an antiseptic (Betadine, Mylan) and topical anesthetic (Xylocaine, Aspen). The head bar, which contained a circular opening, was positioned over the posterior left hemisphere and provisionally fixed with glue (Pattex Sekundenkleber, Henkel). The head bar was then firmly attached to the skull with C&B Superbond (Sun Medical). A circular craniotomy slightly exceeding 4 mm in diameter was made using a dental drill. This craniotomy exposed the left visual cortex. A single-layer circular 4 mm-diameter glass cranial window was placed inside the craniotomy, fixed with glue and stabilized by dental cement (Simplex Rapid, Kemdent). A small metal ring (diameter: 1.8 cm, height: 3 mm) was fixed to the head bar with dental cement for light shielding. Finally, all exposed parts of the skull were sealed with dental cement. Mice were provided with analgesia and antibiotics by default by mixing carprofen and enrofloxacin (Baytril, Bayer) into the drinking water for at least three days after surgery.

### Binocular setup

We designed and built a custom setup to stimulate the eyes of mice independently. Two projectors (ML1050ST, Optoma) were mounted on a custom table. Cut-outs of high-contrast linear polarizing film (XP42-18, Edmund Optics) were fixed in front of the projector lenses and rotated so that their polarization axes were orthogonal to each other. The projectors were aimed at an acrylic glass polarization-preserving screen (ST-Pro-X, Screen Tech-Shop). We built mouse glasses using custom 3D printed material and the same polarizing film as above. The polarization film’s rotation for each eye was adjusted such that its polarization axis aligned with the axis of one of the projectors’ filters, blocking out the light from the other projector (Supplementary Figure 5A). The mouse glasses were held by a mechanical arm (LC mini holder LC6200, Noga) and positioned in front of the mouse using a spirit level to maintain polarization axis alignment. Extinction was verified using camera recordings and a custom-built photometer.

### Microscope

GCaMP6f fluorescence was imaged using a Leica DM6000B microscope. For two-photon imaging, GCaMP6f proteins were excited using a Spectra-Physics Mai Tai mode-locked Ti:sapphire laser with a wavelength of 920 nm. A plane of 570 x 570 μm at a depth of 100 – 200 μm was imaged through a 16x objective (Leica HC FLUOTAR L 16x/0,60 IMM CORR VISIR) using a Leica SP5 resonant mirror scanner operating at a sampling frequency of 27.5 Hz (bi-directional scanning). For previous studies using the same setup, see (Goltstein et al., 2015; Montijn et al., 2016). For widefield imaging, which was used for retinotopic mapping, we recorded the brain surface under green LED light (470 nm; LED: Thorlabs M470L5, driver: Thorlabs T-Cube) through the cranial window using a 1.25x objective (Leica HC PL FLUOTAR 1,25x/0,04 T), filter cube (Leica I3 DM 513828) and a camera (Basler ace 2 a2A1920-160 um).

### Retinotopic mapping

Before two-photon imaging, we charted the visual areas inside the cranial window by analyzing the widefield calcium responses to moving bar stimuli. We presented anesthetized mice (<1% isoflurane, 5 mg / kg xylazine SC) with flashing checkerboard moving bar stimuli on their right eye as described in (Zhuang et al., 2017). More details on the anesthesia can be found in the corresponding section on two-photon imaging below. We showed bar stimuli of four different movement directions with 20 repeats per direction. We analyzed widefield calcium data using custom Python 3 code based on similar analysis by (Zhuang et al., 2017). Briefly, we divided the widefield videos into trial clips and averaged trials with the same bar movement direction. We obtained a phase map for each movement direction through Fourier transform of the averaged videos at the bar sweep frequency. We combined phase maps of opposing directions into azimuth and altitude maps after smoothing with a gaussian 2D filter. Finally, we computed gradient maps of the azimuth and altitude maps and combined them into a single sign field map.

### Two-photon imaging

#### Wakefulness

Awake mice were habituated to sitting in a cylindrical tube while being head-fixed (Goltstein et al., 2015; Montijn et al., 2016). For head fixation, their head bar was attached to a custom tiltable head bar holder using screws. For two-photon imaging, the cranial window was cleaned with distilled water and then covered with a mixture of water and ultrasound gel. The head bar holder with the mouse was then placed on the microscope stage. To avoid rotation of the binocular field of view in comparison to the screen, the angle of the head bar was adjusted by tilting the head bar holder such that the mouse’s eyes were approximately level to the ground. To prevent stimulus light from entering the objective, a custom 3D printed shielding tube was attached to the objective on one end and to a metal shielding ring that was attached to the head bar on the other end. Two cameras (Basler ace acA1300-200 um, Basler) together with custom infrared lights were set up to record the eye movements of the mouse. The mouse glasses were positioned in front of the mouse’s eyes using a mechanical arm (see also section on binocular setup). Then the binocular setup was positioned centrally in front of the mouse and stimuli were presented while two-photon activity was recorded.

#### Anesthesia

Anesthetized recordings were similar to awake recordings except that anesthesia was induced at 4 – 5% isoflurane in 100% oxygen before head fixation. Mice were then head fixed as in awake recordings and a constant supply of anesthetic was provided through a custom-made isoflurane delivery system (Goltstein et al., 2015). We injected mice with xylazine (5 mg / kg s.c.; Sedaxylan, Dechra) after induction to prevent eye drift that may occur under low isoflurane anesthesia in rodents (Nair et al., 2011). See-through eye drops (Lubrithal, Dechra) were applied to keep eyes moist during the recording. Body temperature was controlled using a heating pad, temperature probe and temperature controller (TCAT-2LV, Physitemp instruments). We monitored heart and breathing rate using a custom-built piezo-based device. After xylazine injection, isoflurane levels were lowered to 0.25 – 1%. For each recording, we adjusted the isoflurane level within this range such that we could see stimulus-evoked activity during live imaging without movement of the mouse and with a stable breathing rate below 130 bpm. An example image of an anesthetized mouse in the recording setup is shown in Supplementary Figure 5B. Since no effort was made to find similar imaging planes during awake and anesthetized recordings, there was likely no or little overlap between the target-selective cells in the two states.

### Visual stimulation

All stimuli were generated with custom Python 3 code using the *PsychoPy* package (Peirce et al., 2019). Mice were presented with two different stimulation paradigms that we refer to as experiment 1 (simultaneous-onset binocular conflict) and experiment 2 (delayed-onset binocular conflict). In all experiments, the target consisted of a circular monocular drifting grating (spatial frequency: 0.1 cpd, temporal frequency: 2 Hz) presented at 30 visual degrees size. The mask consisted of a square overlay of 50 pseudorandomly positioned, sized and colored grayscale rectangles that spanned 35 visual degrees. The rectangle side lengths were constrained to 5 - 15 visual degrees length. The rectangle pattern was changed (“flashed”) every 100 ms. The resulting flashing frequency (10 Hz) was below the critical flicker frequency of mice which is estimated at 25 Hz under low-light conditions (Umino et al., 2018). The mask pattern sequence was generated at the start of a recording and stayed the same within that recording. Both target and mask were presented at the horizontal center of the binocular field of view and at 10 degrees elevation. All stimuli were presented on a grey background and were preceded by an intertrial interval of 1.5 – 2.5s consisting only of the background. Each stimulus configuration was shown 20 times per session.

In experiment 1, all stimuli were shown for 1s. We presented three stimulus conditions: (1) target-only (TO), which consisted of one of eight monocular drifting gratings (four orientations x two eyes; 0, 90, 180 or 270 degree orientation); (2) mask-only (MO) which consisted of one of two monocular flashing Mondrian patterns (i.e. either on the left or on the right eye) and (3) binocular conflict (BC), which consisted of a target on one eye and a mask on the other eye. In total, we presented 18 different stimulus configurations in experiment 1 (eight TO configurations, two MO configurations, eight BC configurations).

In experiment 2, stimuli were shown for 6s. In the main text, we refer to two stimulus conditions: (1) long target-only (LTO) which consisted of one of four monocular drifting gratings (two orientations x two eyes; 0 or 90 deg orientation) and (2) intermittent binocular conflict (IBC) in which a target was shown for 6s on one eye while a flashing achromatic Mondrian mask appeared 2s after target onset for 2s on the other eye. This amounts to eight stimulus configurations (four LTO configurations, four IBC configurations). In the supplement, we refer to a third condition: (3) target-only reappear (TOR) in which a monocular target was shown for 2s, then removed for 2s, then represented for 2s.

### Analysis of two-photon imaging data

All analysis of two-photon imaging data was done using custom code in Python 3. We highlight relevant packages, functions and literature where they apply.

#### Preprocessing

A first motion correction was done using the CaImAn toolbox (Giovannucci et al., 2019). A custom-made algorithm removed bidirectional scanning artefacts. Further motion correction and source extraction was done with Suite2p (Pachitariu et al., 2017). From the source extraction, we only included regions-of-interest (ROIs) that were classified as cells using Suite2p’s built-in classifier with a probability greater than 50%.

We computed baseline-corrected fluorescence values (dF/F) as described by (Chen et al., 2013). For each fluorescence trace, we subtracted neuropil activity, then scaled the fluorescence value of each trace in each trial based on its average fluorescence during a 1s prestimulus period.

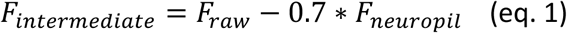

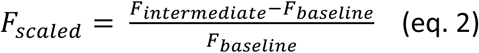

Finally, we smoothed the scaled fluorescence values with a gaussian kernel (sigma = 50 ms) and upsampled them to 100 Hz before further analysis.

#### Brain area determination

Brain areas were determined by manually overlaying the brain surface two-photon image of a recording site over the widefield anatomical image and retinotopic map. If a recording site was found to contain multiple brain areas, custom software was used to attribute brain areas to each suite2p ROI. Only cells in brain areas V1 and LM were included in the analysis.

#### Target-selective cells

Target-selective cells were identified according to three criteria. (1) *Selective responsiveness*: for each cell, a Kruskal-Wallis test was conducted to test for differences between responses to target stimuli. There were eight target stimuli in experiment 1 and four target stimuli in experiment 2; each target stimulus was shown at least 20 times. In experiment 1, the average activity during 1s stimulation was taken for each trial. In experiment 2, the average activity during the first 2s of stimulation were taken for each trial. P-values were corrected for every experiment and stated separately using FDR correction (Benjamini Hochberg procedure) and only cells with p_adjust_ < 0.05 were included. This step excluded cells that did not respond reliably to any target stimulus or that responded similarly to all target stimuli. (2) *Strong responsiveness to preferred target*: The preferred target stimulus was determined by selecting the target stimulus that elicited the maximal average response r_pref_. Only cells with r_pref_ > 0.25 dF/F were included. This excluded cells that only weakly responded to their preferred stimulus. (3) *Low baseline variance:* for each cell, we computed the standard deviation across all trials for 1s before stimulus onset σ_baseline_. We only included cells with σ_baseline_< 1 dF/F. This excluded cells with unacceptable noise levels, for example those with very low baseline fluorescence for which dF/F values were unreliable.

#### Mask-selective cells

Mask-selective cells were identified similarly to target-selective cells, except now only two mask stimuli were available (left eye, right eye). For the first criterion, the Kruskal-Wallis test was therefore replaced by a two-sided Mann-Whitney U test. All other criteria remained unchanged.

#### Trial selection

For further analysis, for each cell, only trials that included the preferred stimulus were included in the analysis. That is, for target-selective cells in experiment 1, only trials where the TO condition consisted of the preferred target and trials where the BC condition consisted of the preferred target in conflict with a mask were included. For experiment 2, the same applies for LTO and IBC population responses. For mask-selective cells (Figure 6), only trials where the MO condition consisted of the preferred mask were included and similarly only BC trials which consisted of the preferred mask and a target on the other eye were included.

#### Population responses

Population responses (e.g. Figure 2C) for each stimulus condition were computed in two steps (nested averaging). First, for each cell, the average response over the 20 repeats of that condition was computed. Second, an average of the cell averages was computed. Shaded areas around population response curves represent the standard error of the 2nd mean which is the standard deviation across cells divided by the square root of the cell count.

#### Effective differences

Effective differences between condition responses were assessed by computing Cohen’s d. Cohen’s d is defined as

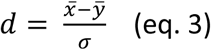

where 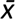 and 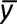 are the means of two sets of values *x* and *y,* and *σ* is the pooled standard deviation of the two groups.

To quantify the differences between TO and BC conditions in experiment 1 in a single value (e.g. Figure 2C), for each cell, the average activity during 1s of stimulation across all repeats was computed. These averages were arranged in a vector for each condition separately. Then, Cohen’s d was computed between the vectors of TO and BC responses. The statistical significance of differences between population responses was assessed through two-sided Wilcoxon rank sum tests using the same vectors. In experiment 2, the same steps were performed for LTO and IBC trials with the only difference being that the effective difference was computed for two periods: (1) the period from t=3s to t=4s after stimulus onset which captured binocular conflict responses in IBC trials but excluded the transient of those responses at mask onset; (2) the period from t=5s to t=6s which captured the responses after mask offset in IBC trials but again excluded transient responses. To quantify the effective differences between two conditions continuously over time (e.g. Figure 3D), the same procedure was used except the average activity was computed using a sliding window of 250 ms.

#### Suppression of individual cells

For each cell in experiment 1, a suppression score *s* was calculated as follows:

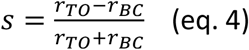

Here, *r_BC_* is the average activity of a cell during 1s of stimulation in BC trials and *r_TO_* is similar but for TO trials. Because of criterion (2) for target-selective cells, *r_TO_* could not be smaller than 0.25 dF/F. *r_BC_* values that were smaller than 0 were set to 0 such that the suppression scores were limited to [-1, 1]. A suppression score of 1 indicated that a cell was only active in the TO but not active in the BC condition. Significance of response modulations in experiment 1 was assessed for each cell by a two-sided Mann Whitney U test between average responses (1s) during TO and BC trials (20 repeats per condition). P-values were corrected using FDR correction (Benjamini Hochberg procedure) separately for each state. Cells with p_adjust_ >= 0.05 were referred to as unaffected. Cells with p_adjust_ < 0.05 and s > 0 were referred to as suppressed, while cells with p_adjust_ < 0.05 and s < 0 were referred to as enhanced. In experiment 2, suppression scores were calculated similarly to experiment 1 but now for LTO and IBC trials. Akin to the effect size calculation of experiment 2, suppression scores were calculated separately for time periods t=3 - 4s and t=5 – 6s after stimulus onset.

#### Decoding

We decoded target orientation from neuronal responses in the awake simultaneous-onset experiment, for each recording (n = 45) separately, using linear support vector machines. There were four possible target orientations. For this, we averaged the response of each neuron during the 1s stimulation period of each trial and then arranged the neuronal responses in a matrix with shape (# trials, # neurons). We split the trials by two factors: the first factor was the trial type (target-only, binocular conflict). The second factor was the target eye (any, contralateral, ipsilateral). In each split, there were at least 20 trials of each target orientation. For evaluation, we used stratified k-fold cross-validation (k = 10) and quantified performance as the average balanced accuracy score from the ten folds.

#### Binocular rivalry attractor model

To understand the results within the broader context of the primate BR literature (which are canonically modelled with attractor networks), we fit the BR model of (Wilson, 2007) to the neural data from experiment 1. The model consists of a system of four nonlinear differential equations governed by competitive inhibition between target- and mask-driven populations that slowly reduce their firing rates due to spike-dependent adaptation.

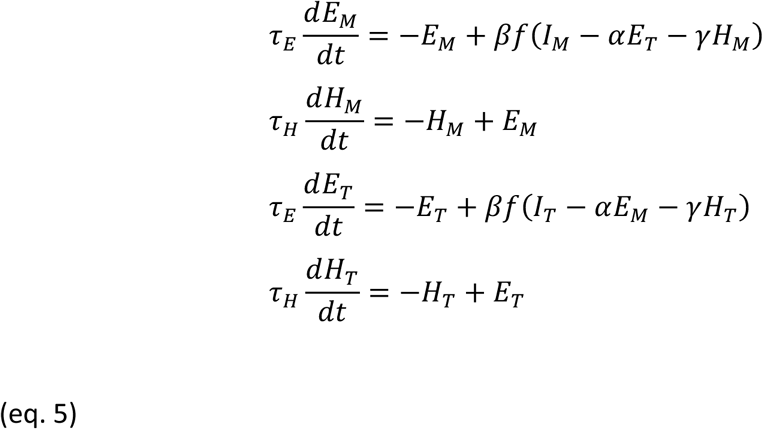

The four state variables (𝐸_𝑀_, 𝐸_𝑇_, 𝐻_𝑀_, 𝐻_𝑇_) represent the average neural activity of populations driven by the mask (𝐸_𝑀_) and target stimulus (𝐸_𝑇_), and the average slow self-inhibitory adaptation current of the mask (𝐻_𝑀_) and target (𝐻_𝑇_) populations. The transfer function 𝑓 is a simple threshold linear function 𝑓(𝑥) = max (𝑥, 0). 𝐼_𝑀_and 𝐼_𝑇_are dimensionless parameters representing the strength of the mask and target inputs. The parameters 𝛼, 𝛽 and 𝛾 are dimensionless. 𝛼 represents the strength of the competitive inhibition.

𝛽 represents the gain of the population transfer function. 𝛾 represents the strength of spike-dependent adaptation. The timescale parameters 𝜏_𝐸_and 𝜏_𝐻_are in units of ms.

The model was integrated numerically using the Euler-Maruyama method with a timestep of 1 ms (smaller time steps did not alter results) with weak additive noise (𝜎 = 0.001) in the neural variables 𝐸_𝑀_ and 𝐸_𝑇_,.

#### Parameter determination

We determined the value of 𝜏_𝐸_by fitting a decaying exponential (𝑎𝑒^−𝑡/τ^ + 𝑏) to the neural data from experiment 1 and taking the average of 𝜏 across awake conditions. The value of 𝜏_𝐻_ was constrained by prior literature (Moreno-Bote et al., 2007; Wilson, 2007). We fit the model via a four-step grid search to find the parameters that best minimized the mean squared error (MSE) between the simulated neural activity (averaged across 25 trials) and the empirical data. We started by fitting 𝛽and, 𝛾 and 𝐼_𝑇_ to the awake TO condition. In this step, 𝛼 and 𝐼_𝑀_ could be ignored as competitive inhibition does not play a role in the dynamics in the absence of the mask. We then fitted 𝛼 and 𝐼_𝑀_ for the BC condition (fixing the parameters found in the TO condition). We repeated this procedure for the anesthetic conditions taking the target and mask strength parameters found in the awake condition as fixed (as these features are external to the effects of anesthesia). Fitted parameter values and the range of the grid search are shown in the following table:

**Table 1.**
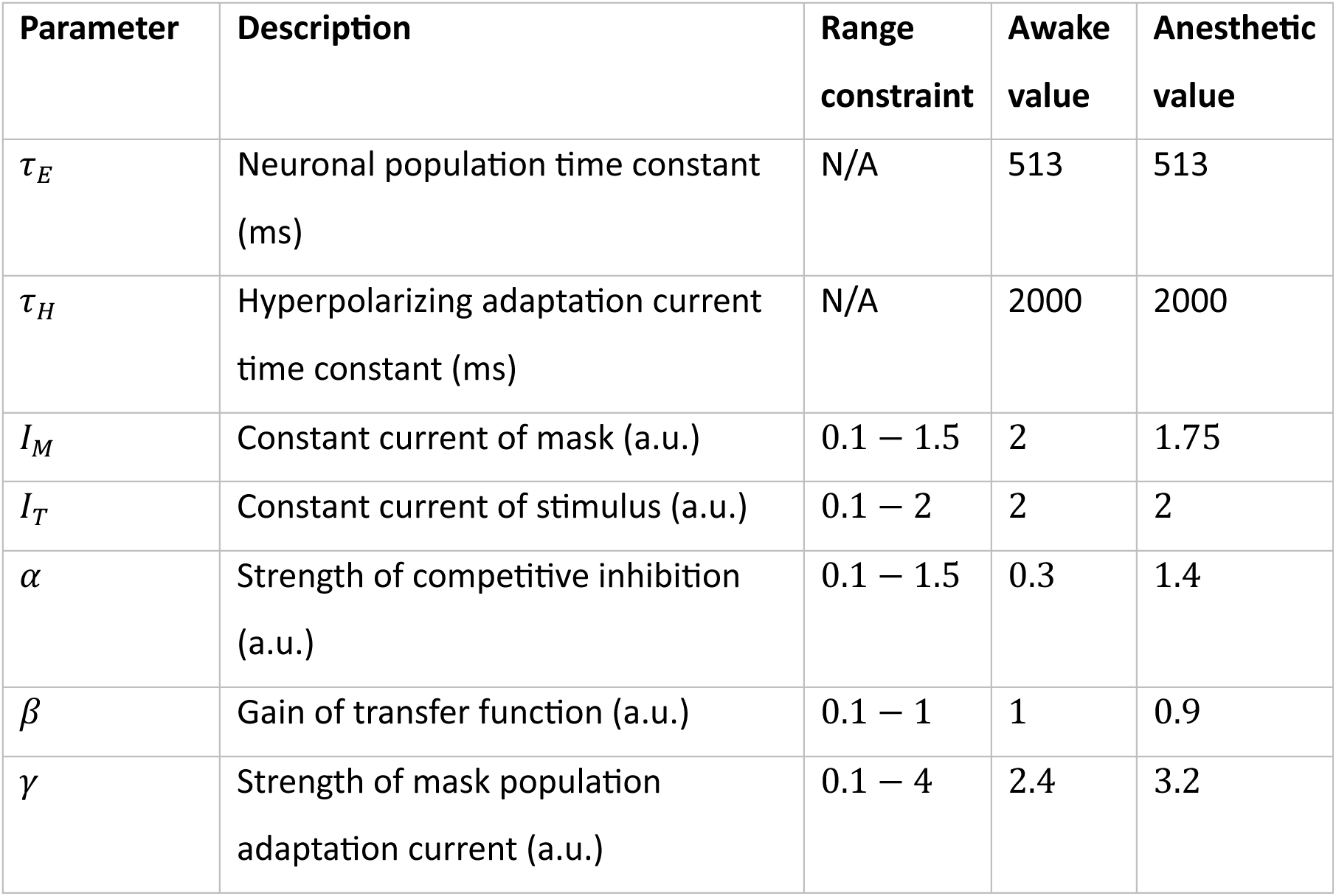
Modeling parameter values.

| Parameter | Description | Range constraint | Awake value | Anesthetic value |
| --- | --- | --- | --- | --- |
| $\tau_E$ | Neuronal population time constant (ms) | N/A | 513 | 513 |
| $\tau_H$ | Hyperpolarizing adaptation current time constant (ms) | N/A | 2000 | 2000 |
| $I_M$ | Constant current of mask (a.u.) | 0.1 – 1.5 | 2 | 1.75 |
| $I_T$ | Constant current of stimulus (a.u.) | 0.1 – 2 | 2 | 2 |
| $\alpha$ | Strength of competitive inhibition (a.u.) | 0.1 – 1.5 | 0.3 | 1.4 |
| $\beta$ | Gain of transfer function (a.u.) | 0.1 – 1 | 1 | 0.9 |
| $\gamma$ | Strength of mask population adaptation current (a.u.) | 0.1 – 4 | 2.4 | 3.2 |

## Author contributions

MB: Conceptualization, Data curation, Formal analysis, Investigation, Methodology, Software, Validation, Visualization, Writing

LE: Methodology, Resources CW: Software

GH: Methodology

MS: Resources, Supervision, Writing

CP: Conceptualization, Funding acquisition, Methodology, Resources, Supervision, Writing

