## Supplemental Figures for "Continuous Flash Suppression responses in mouse visual cortex: stimulus laterality and anesthesia effects"

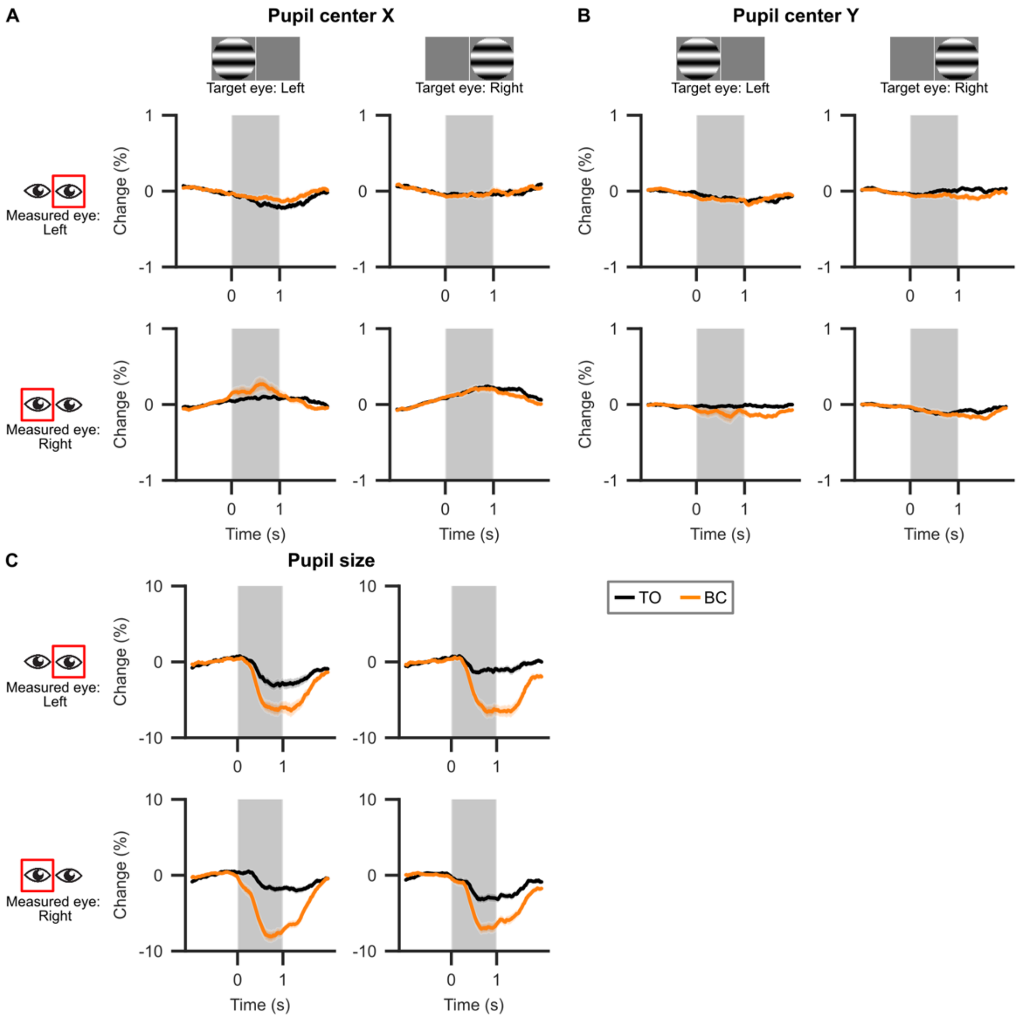


**Supplementary figure 1. Pupil measurements during awake simultaneous-onset experiment.**
(A) Change of x-position of pupil center during stimulus presentation compared to x-position during baseline (1s before stimulus onset). Changes are plotted separately for target displayed on the left or right eye (columns) as well for left and right pupil (rows). Averaged over n = 34 recordings (from seven out of eight mice) Shaded region around lines indicates standard error of the mean. Shaded grey region indicates stimulus presentation period. TO = target-only, BC = binocular conflict.
(B) Like (A) but now for Y-position of pupil center.
(C) Like (A) but now for pupil size.
Overall, the pupils of mice did not change positions during stimulus presentation systematically as all average position changes were smaller than 0.5%. We saw systematic pupil constriction upon stimulus onset which was higher for the BC condition but similar for both eyes. Therefore, we conclude that the observed differences of contra- and ipsi-preferring cell populations did not arise from differential eye movements or pupil constriction.

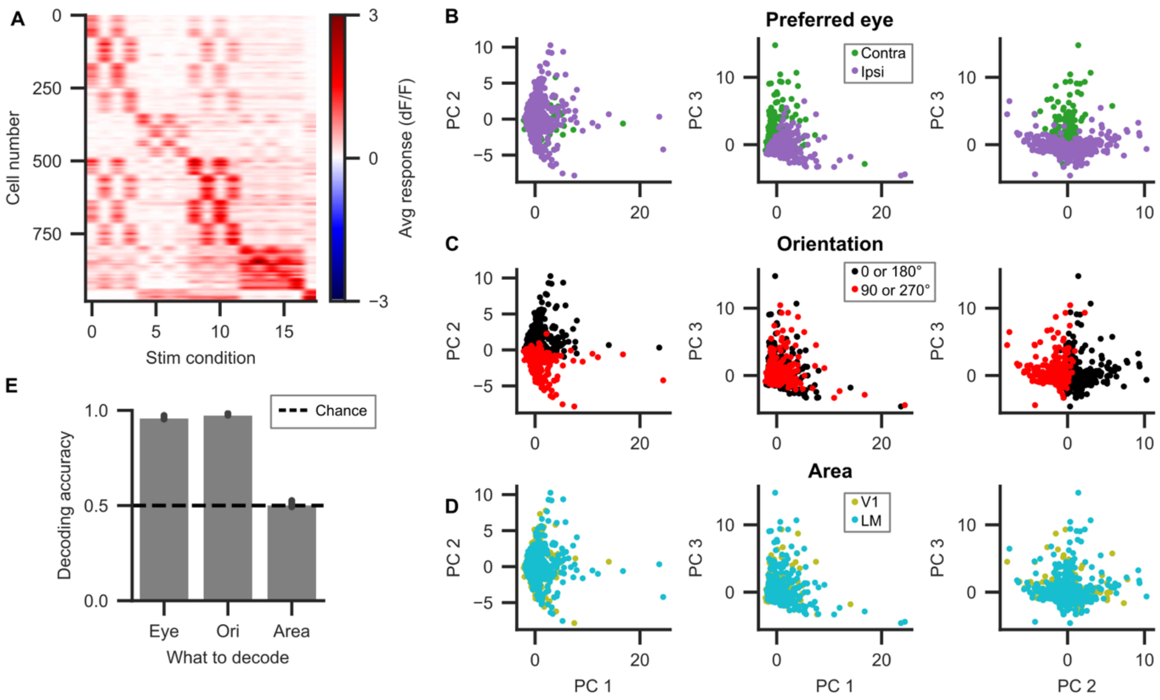


**Supplementary figure 2. V1 and LM cells are indistinguishable by linear classification.**
(A) Average responses of all cells in the awake simultaneous-onset experiment to all 18 stimulus conditions (eight monocular targets, eight binocular conflict displays, two monocular masks). Cells were sorted by condition with highest response for visualization purposes.
(B) Scatterplots of neuronal response patterns (i.e. the average responses during stimulus presentation for all 18 stimulus conditions) after transformation by PCA. Each dot represents a cell. Dots are colored based on the eye of origin of the preferred target. PC = Principal component. Contra = preferred target on contralateral eye, Ipsi = preferred target on ipsilateral eye.
(C) Same as (B) but now colored by preferred orientation.
(D) Same as (B) but now colored by brain area.
(E) Accuracy of decoding cell characteristics from their average responses to all stimulus conditions (depicted in A) using a linear support vector machine classifier. Eye = Eye of origin of preferred target (depicted in B). Ori = Orientation of preferred target (depicted in C). Area = Brain area (depicted in D). Dotted line indicates chance level decoding (50%).
Whereas response properties such as preferred orientation and preferred eye could be reliably decoded from the cell response patterns, whether a cell was located in V1 or LM could not be decoded above chance level. This indicates that V1 and LM cells had highly similar response patterns in our data. Therefore, we decided to pool V1 and LM cells in this study.

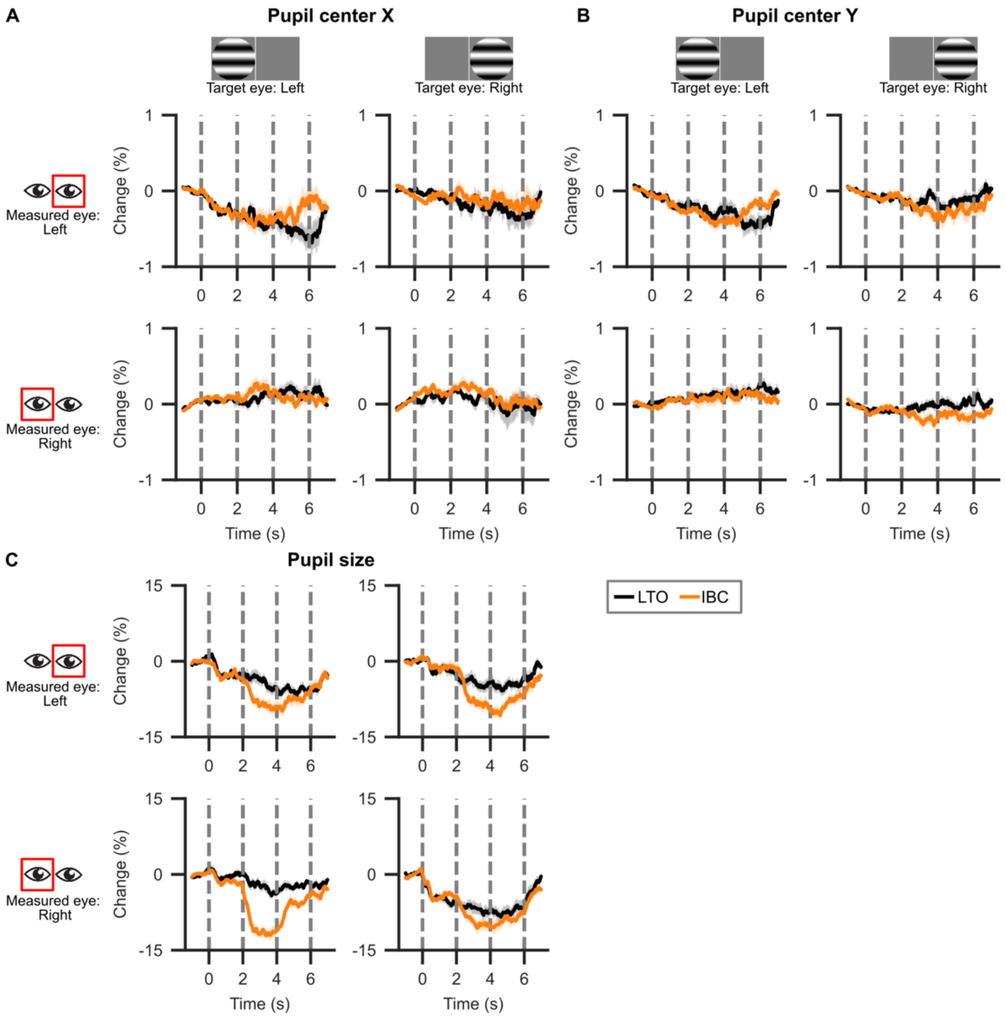


**Supplementary figure 3. Pupil measurements during awake delayed-onset experiment.**
(A) Change of x-position of pupil center during stimulus presentation compared to x-position during baseline period (1s before stimulus onset). Changes are plotted separately for target displayed on the left or right eye (columns) as well for left and right pupil (rows). Averaged over n = 16 recordings from four mice. Shaded region around lines indicates standard error of the mean. Dashed grey lines indicate stimulus timings (target onset, mask onset, mask offset, target offset). LTO = long target-only, IBC = intermittent binocular conflict.
(B) Like (A) but now for Y-position of pupil center.
(C) Like (A) but now for pupil size.
As in Supplementary figure 1, the average pupil position changed less than 0.5% compared to its base position during stimulus presentation. While pupils were more constricted during IBC than LTO trials, pupil constrictions were similar across eyes. From this, we conclude that differences between contra- and ipsi-preferring cell populations did not arise from differential eye movements in the delayed-onset experiment.

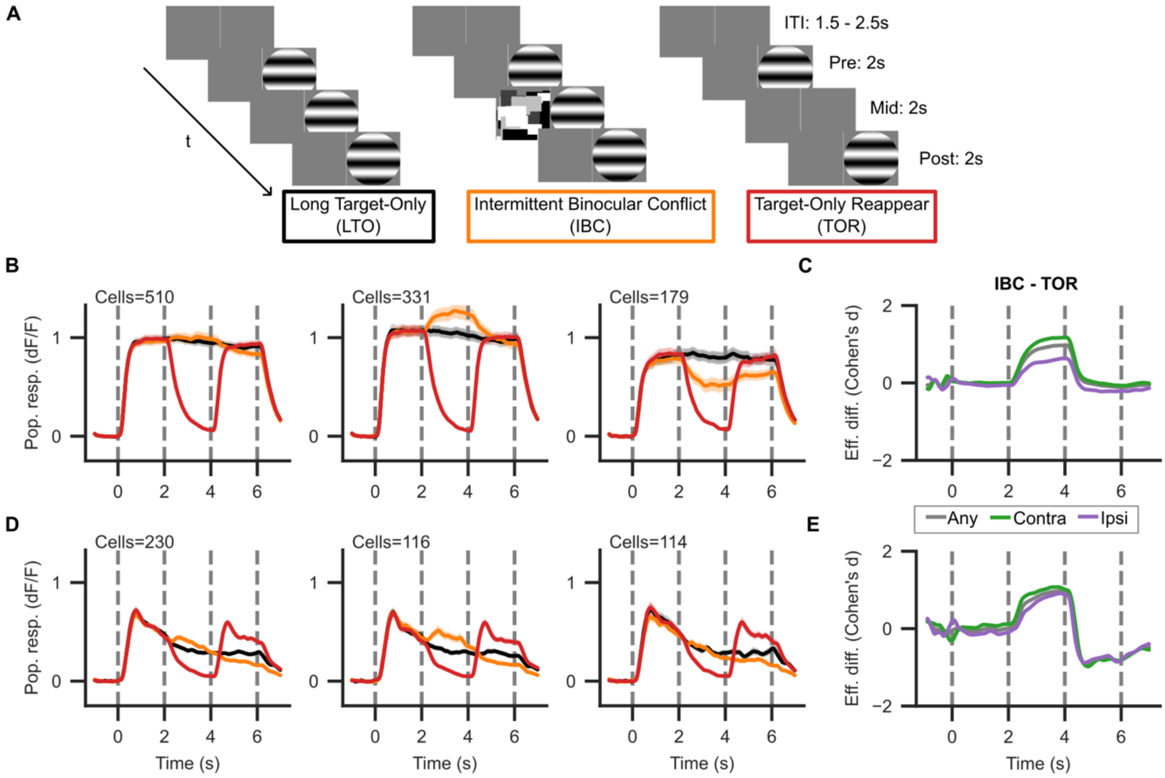


**Supplementary figure 4. Population responses in delayed-onset paradigm with an extra control condition.**

(A) Stimulus paradigm as in Figure 5A but now including an extra control condition: target-only reappear (TOR).

(B) Population responses as in Figure 5D but now including the TOR control condition. The data for LTO and IBC conditions are the same as in Fig.5.

(C) Effective difference (Cohen's d) between IBC and TOR population responses.

(D) Same as B but now under anesthesia.

(E) Same as C but now under anesthesia.

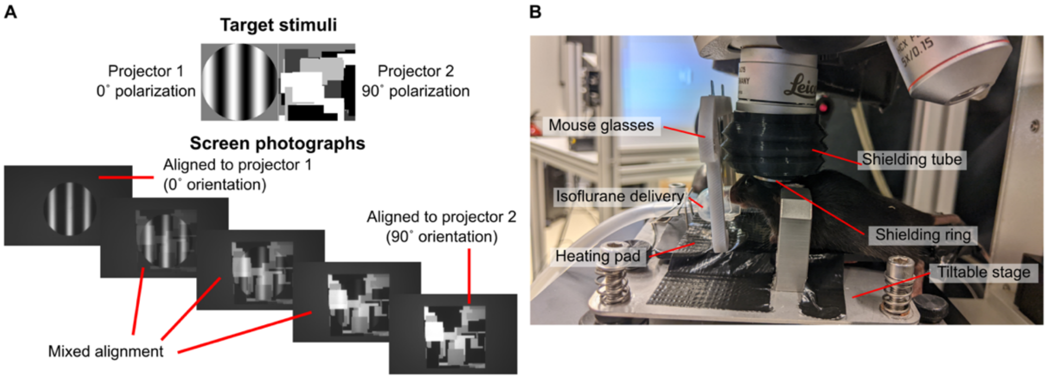


**Supplementary Figure 5. Stimulus and recording specifics.**
(A) Demonstration of exclusion through polarization filters. (Top) Schematic representation of binocular stimuli. (Bottom) Photographs through rotating polarization filter to show polarizer alignment.
(B) Photo of anesthetized mouse under microscope with annotated experimental elements.
